# Self-Assembly of Tunable Intrinsically Disordered Peptide Amphiphiles

**DOI:** 10.1101/2022.10.28.514176

**Authors:** Tamara Ehm, Hila Shinar, Guy Jacoby, Sagi Meir, Gil Koren, Merav Segal Asher, Joanna Korpanty, Matthew P. Thompson, Nathan C. Gianneschi, Michael M. Kozlov, Salome Emma Azoulay-Ginsburg, Roey J. Amir, Joachim O. Rädler, Roy Beck

## Abstract

Intrinsically disordered peptide amphiphiles (IDPAs) present a novel class of synthetic conjugates that consist of short hydrophilic polypeptides anchored to hydrocarbon chains. These hybrid polymer-lipid block constructs spontaneously self-assemble into dispersed nanoscopic aggregates or ordered mesophases in aqueous solution due to hydrophobic interactions.

Yet, the possible sequence variations and their influence on the self-assembly structures is vast and have hardly been explored.

Here, we measure the nanoscopic self-assembled structures of four IDPA systems that differ by their amino acid sequence. We show that permutations in the charge pattern along the sequence remarkably alter the headgroup conformation and consequently alters the pH-triggered phase transitions between spherical, cylindrical micelles and hexagonal condensed phases. We demonstrate that even a single amino acid mutation is sufficient to tune structural transitions in the condensed IDPA mesophases, while peptide conformations remain unfolded and disordered. Furthermore, alteration of the peptide sequence can render IDPAs to become susceptible to enzymatic cleavage and induces enzymatically activated phase transitions.

These results hold great potential for embedding multiple functionalities into lipid nanoparticle delivery systems by incorporating IDPAs with desired properties.

## Introduction

Self-assembly of amphiphiles that combine hydrophilic and hydrophobic molecular moieties plays an omnipresent role both in natural and synthetic systems. In the biological world, lipid self-organization lies at the basis of cell membrane integrity, transport vehicles, and reaction vessels with precisely controlled size and functionality. In pharmacology, synthetic amphiphiles, in addition to natural lipids, are used to form nanoscopic carriers for encapsulating drugs.^1^ Following rational design principles, control of size and stability of assemblies is achieved most prominently by using polyethylene glycol (PEG)-lipid conjugates. These strategies result in highly efficient formulations such as lipid nanoparticles that serve as RNA-based vaccine carriers against SARS-CoV-2,^2^ or other cargos or drugs.^3–9^ In order to advance functionality, nanocarriers composed of stimuli-responsive (e.g., enzymatic, pH, temperature) amphiphilic systems^10–16^ are studied, as they can potentially reduce the side effect of drugs by targeted release in tissues.

Amphiphiles can self-assemble into various mesophases in solution. Their mesoscopic morphology is, to a first approximation, determined by the volumetric ratio of the effective hydrophilic head group to the hydrophobic tail, as described by the so-called packing parameter.^17^ Here, the hydrophobic domain is composed of one or two fatty acid-based chains, as we previously demonstrated.^18^ In recent works, polypeptide chains have been conjugated to a hydrophobic domain to create peptide amphiphiles.^19–22^ In these studies, the polypeptides exhibited folded conformations and formed well-controlled nanoscale assemblies, such as long nanorods, that proved capable of encapsulating and releasing small molecules.^23,24^ The folded hydrophilic headgroup can lead to specific and relatively rigid structures that specific enzymes can recognize. Thus, these structures are potentially beneficial in applications where specific ligand-receptor binding is required.^25,26^

As in many other cases in biology, liquid-like structures dominated by weak and reversible interactions can be leveraged for novel biomedical applications. Indeed, and in contrast to the central dogma of proteins’ folding, about half of the proteome contain proteins, and large domains that do not fold into rigid secondary or tertiary structures.^27–29^ These unfolded, intrinsically disordered proteins (IDPs) provide a significant functional advantage, enabling them to interact weakly with a broad range of binding partners, including themselves.^30,31^ Prominent examples of IDPs with weak interactions (i.e., on the order of thermal energy) include IDPs occurring in liquid-liquid phase separations^32^ or forming selective filters in nucleoporin complexes.^33^ Other examples of long disordered domains are the carboxy tails of intermediate filaments proteins. These proteins retain their disordered nature, even when constrained at high-density^31,34,35^ and are responsible for fine-tuning the mechanical cytoskeleton behavior.^36–40^

Previous works showed that both the sequence composition and the fraction of charged amino-acids play essential roles in the properties of a protein’s unfolded ensemble.^41,42^ For example, molecular dynamic simulations suggest that sequence composition and patterning are well reflected in the global conformational variables (e.g., the radius of gyration and the hydrodynamic radius), but end-to-end distance and dynamics are highly sequence-specific.^43^ Such analysis is suitable for comparing IDPs of different lengths.^29,44^ Moreover, it was demonstrated that the total net charge is inadequate as a descriptor of sequence–ensemble relationships for many IDPs. Instead, sequence-specific distributions of oppositely charged residues are synergistic determinants of conformational properties of polyampholytic IDPs.^45^

Sequence-encoded conformational properties can be extracted by calculating the charge patterning parameter (0 ≤ *κ* ≤ 1) and the fraction of charged residues (*FCR*).^45^ Low values of *κ* point to sequences where intrachain electrostatic repulsions and attractions are balanced. In contrast, high *κ* sequences show a preference for hairpin-like conformations caused by long-range electrostatic attractions induced by conformational fluctuations.^45^ Other studies presented coarse-grain models that identify short-range electrostatic attractive domains between IDPs.^36,37,46^ Altogether, IDPs present an intriguing, unexplored territory that combines the structural plasticity of weakly interacting polymers with the specificity of the amino-acid sequence.

In this context, intrinsically disordered peptide amphiphiles (IDPAs) are of great interest as they combine building blocks from natural lipids and proteins.^47–49^ IDPAs are composed of intrinsically disordered peptides conjugated to hydrocarbon chains, creating amphiphiles with polymeric headgroups and hydrophobic anchors that remain compatible with natural lipid membranes. Though IDPAs hold promise for fine-tuned nanoscopic self-assembly, the sequence space of even a 20 amino-acid short polypeptide is extremely large and hardly explored.

Here, we present an approach to verify that structural transitions in IDPA assemblies depend on the peptide sequence, even though the head group conformation is disordered. We designed IDPAs with a peptide sequence inspired by neurofilament low chain protein and conjugated the sequence to a single or double hydrocarbon tail to compare peptides composed of the same amino acids but in different sequence order. Using small angle X-ray scattering (SAXS) and cryogenic transmission electron microscopy (cryo-TEM), we analyzed the nanoscopic structural phase transitions as a function of pH and buffer salinity. We show that the phase transitions are controlled by the hydrophobic domain and the charge pattern of the peptide sequence that may induce hairpin-like conformations. Surprisingly, although the amphiphiles remain disordered, the mesoscopic structures exhibit low polydispersity. Structural phase transitions in mesoscopic order are sensitive to the mutation of a single amino acid in the polypeptide head group. Finally, we demonstrate that incorporating suitable motifs renders IDPAs enzymatically cleavable. Ultimately, the reported sequence-dependent properties of IDPA mesophases could be exploited for the development of future drug carrier systems.

## Material and Methods

### Synthesis and purification

All peptides were synthesized via solid-phase synthesis and purchased from LifeTein (USA). Amino acids are conjugated from the C-terminus to the N-terminus while the peptide remains anchored to insoluble solid resin support. The process involves repeated coupling cycles, washing, deprotection, and washing. The hydrophobic domain has either single or double hydrocarbon chains. After adding the last amino acid and deprotection, the fatty acid chain was conjugated to the deprotected amine. Double chain PDAs were prepared by conjugation of Fmoc-Lys(Fmoc)-OH, followed by cleavage of the two Fmoc protecting groups and conjugation of the two tails.

### Sample preparation

The IDPA or peptide powder was first fluidized in purified water (Milli-Q) at twice the desired concentration. The solution was then titrated with 1M NaOH to a pH where the solution became more homogeneous (preferably a pH where the IDPAs are soluble in water). Titration was monitored using a pH probe. Following titration, 50 *μ*l of the solution was combined with 50 *μl* of 2X buffer of choice to achieve a pH in the vicinity of the desired one. The 2X buffers Acetic Acid (pH 3-4.5), 2-(N-Morpholino)ethansulfonsäure (MES, pH 5-6.5), and 3-(N-morpholino) propane sulfonic acid (MOPS, pH 7-7.5), were prepared at 200 mM, to achieve final buffer molarity of 100 mM after mixing with IDPA or peptide solution 1:1 (vol:vol).

### CD

Circular dichroism (CD) measurements were performed using a commercially available CD spectrometer (Applied Photophysics Chirascan). IDPs were added to a glass cuvette with a 1*mm* path length. The peptides were mixed with phosphate buffer to achieve a concentration of 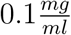. The measurements were performed with phosphate buffer because the buffers used for the X-ray scattering experiments (mainly MOPS and MES) have high absorption at the relevant CD wavelengths. The wavelength range of 190-260 nm was measured in 1-nm steps with 0.5 seconds per point. Three measurements were performed for each sample, and the mean value was calculated.

### Computational methods for disorder analysis

Disorder can also be analyzed computationally. IUPred2^50^ uses an energy estimation method. The principal lies in a 20 × 20 energy predictor matrix *P*_*ij*_ that shows the statistical potential for the 20 amino acid to connect with each other in a globular protein. :

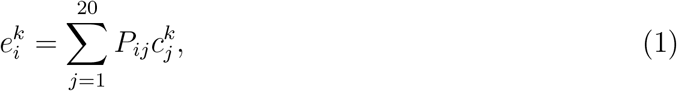

where 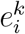 is the energy of the residue in position *k* of type *i*. The equation calculates for each position *k* the sum of for all elements *j* in the amino acid composition vector *c*_*j*_ for all types *i*. The parameters are optimized to minimize the difference between energies estimated from the amino acid composition vector and the energies calculated from the known structure for each residue in the dataset of proteins. As IUPred2, ANCHOR2^50^ also uses an energy estimation method and adds two more terms to the energy estimation: the interaction of the residues with the globular protein and the local environment. Thus, ANCHOR2 combines the disordering tendency calculated by Iurpred with the sensitivity to the environment of the protein and can predict if a specific region is disordered in isolation but can undergo disorder-to-order transition upon binding- without even knowing the possible binding partners. Netsurf 2.0^51^ is a sequence-based method and uses an architecture composed of convolutional and long short-term memory neural networks trained on solved protein structures to predict disorder.

### Cryo-TEM

Cryogenic TEM (cryo-TEM) specimens were prepared using an FEI Vitrobot by blotting in 95% humidity and subsequently plunging lacey carbon grids into liquid ethane. Images were taken for cryo-TEM using a JEOL 1230 transmission electron microscope operating at 120 keV equipped with a Gatan camera.

### FRET

Fluorescence spectra of IDPAs were measured using a Cary Eclipse fluorescence spectrophotometer (Agilent Technologies, Santa Clara, CA). Measurements were done in a 1 cm quartz cuvette at 10 *μ*M concentrations in 100 mM buffer at 25°. Excitation spectra of IDP and IDPA included donor and acceptor (DA) spectra and acceptor only (AO) spectra. The samples were excited over the range of 250–330 nm (bandwidth 2.5 nm), and the emission was set to 350 nm (bandwidth 20.0 nm). The excitation spectra were normalized at 290-295 nm (no Tyr absorption). The level of energy transfer, E, between the donor and the acceptor, Y and W, respectively, was determined by the difference in integrated intensity at 270-285 nm and using YW dipeptide as a reference for 100% energy transfer. Buffer and background signals were routinely measured and subtracted. Distance, r, was calculate using *E* = *R*_0_*/*(*R*_0_ + *r*) while the Forster radius, *R*_0_, was set as 15 *Å*.

### Small angle X-ray scattering (SAXS)

All samples for SAXS were prepared at a final concentration of 5mg/ml, which is an order of magnitude higher than the typical micro-molar CMC of 5*μ*M, reported for similar peptide amphiphiles.^47,48,52,53^ For solubilizing conditions (above the transition pH, generally above pH 6), samples were measured at three synchrotron facilities: Beamline B21, Diamond Light Source, beamline SWING, SOLEIL synchrotron facility, Paris, France, and DESY, Hamburg, Germany. For phase-separating samples that display sediment (below the transition pH, generally pH 3-5.5), measurements were performed using an in-house X-ray scattering system, with a Genix3D (Xenocs) low divergence Cu K_*α*_ radiation source (wavelength of *λ* = 1.54 Å) with a Pilatus 300K (Dectris) detector, as well as beamline I22 at Diamond Light Source. Samples were measured inside 1.5 mm quartz capillaries (Hilgenberg). All 2D measurements were radially integrated using SAXSi^46^ to get 1D Intensity- scattering vector *q* data sets.

### Singular Value Decomposition (SVD)

In SVD, a minimum number of singular vectors represents the entire data set. Thus, these independent curves can represent the entire data set by their linear combinations:

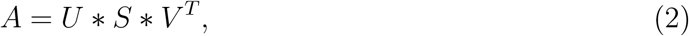

where *U* yields a set of left singular vectors, i.e., orthonormal basic curves *U* (*k*)(*si*), that spans the range of matrix *A*. In contrast, the diagonal of *S* contains their associated singular values in descending order. For our scattering curves, the residuals are calculated via:

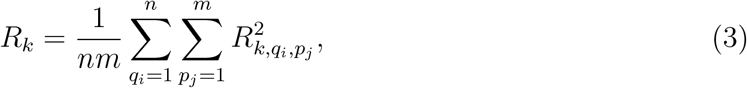

where *m* is the size of the scattering vector *q* and *n* are the number of pH steps, are plotted as a function of the number of singular vector components (*k*) that were chosen to reconstruct the data matrix. 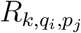 is defined by 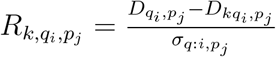, where *D* is the data matrix, in which each column represents a one dimensional scattering curve, *I*(*q, p*) at every pH step *p. D*_*k*_ is the reconstructed data matrix using *k* singular orthonormal vectors, and each term (*q*_*i*_, *p*_*j*_) in the matrix *σ* corresponds to the measured standard error for the corresponding term in *D*.

## Results

### IDPA Primary Structure

In the presented study, all IDPs are directly conjugated to fatty acids of various lengths to create the amphiphilic IDPAs. This study used various standard linear fatty acid chains with 12 (Lauric acid), 14 (Myristic acid), 16 (Palmitic acid), and 18 (Stearic acid) carbons (table 1 for crucial parameters of IDPAs and Fig. 1 for chemical structures). The IDPAs were synthesized using an automated solid-phase synthesizer. Thus, the molecular architectures are highly tunable, allowing us to study various hydrophobic and hydrophilic domains in a controlled manner. The peptide sequences are 18 amino acids long, containing protonable residues and hydrophilic amino acids (Supplementary Fig. S.1).

**Table 1:**
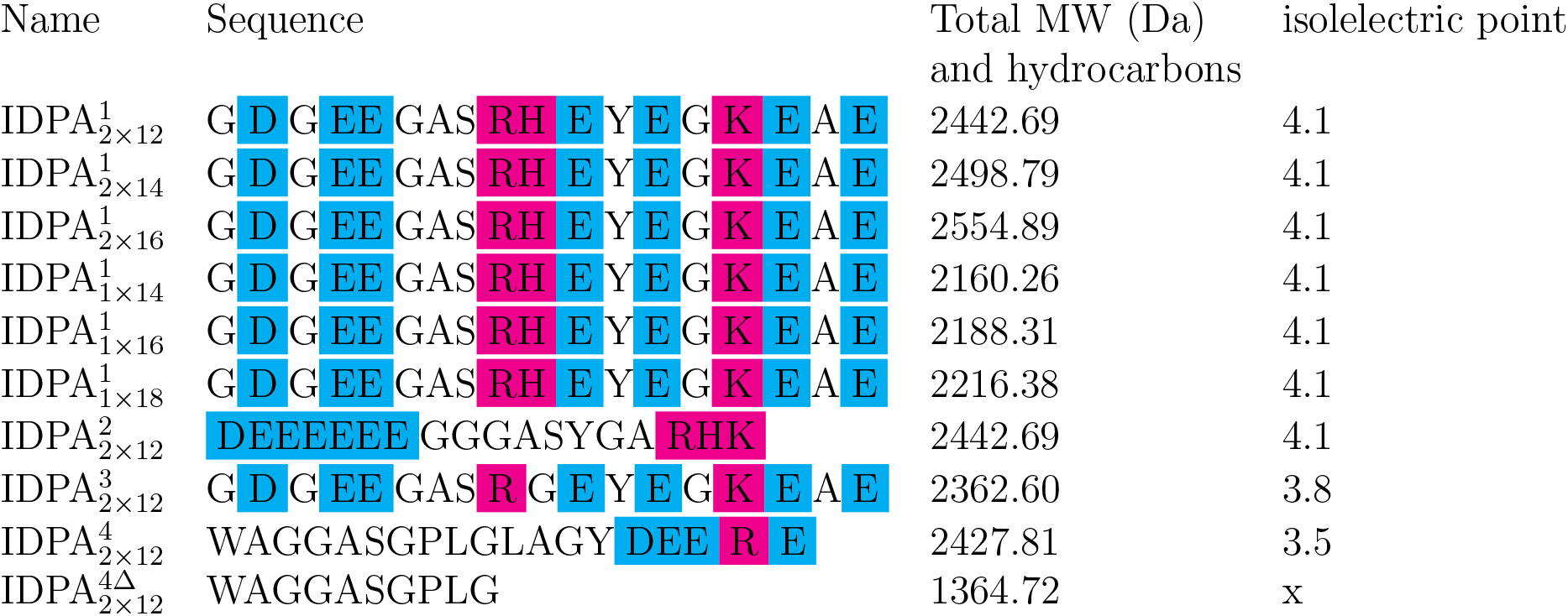
Key paramteres and notation for IDPAs used in this paper. Blue colored letters stand for anionic amino acids, pink colored ones for cationic amino acids. Upper case number in IDPA name is the sequence number and lower case numbers are the number of tails in this molecule.

**Figure 1:**
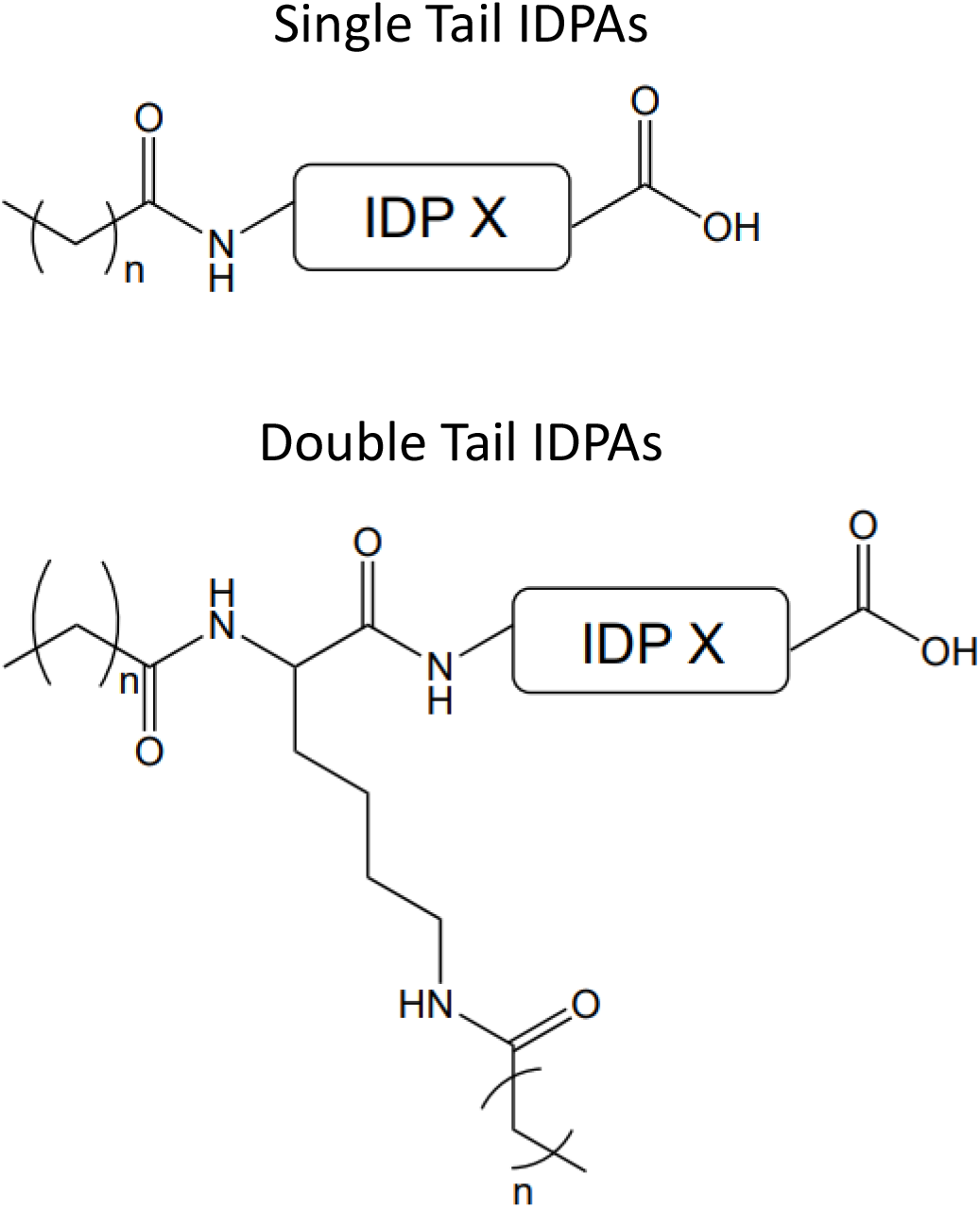
Chemical structures for double and single tailed IDPAs. For detailed chemical structures of IDP X see Supplementary Fig. S.1.

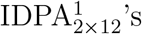 primary sequence (Table 1, Supplementary Fig. S.1) is inspired by the intrinsically disordered carboxy tail-domain of neurofilament-light (NF-L) protein found in the cytoskeleton of nerve cells.^35–37,54^ In previous research,^47^ we introduced this IDP sequence to create IDPAs where aromatic branchings units were used to cap the N-terminus of the IDP sequence and allow the branching into two different types of architectures containing either two or four hydrocarbon tails (2×12, 4×7). We showed that these IDPAs undergo a sharp phase transition from low-dispersity micellar spheres to extremely elongated wormlike micelles. Here, we present IDPAs that can be entirely prepared using conventional solid phase peptide synthesis and allow us to study alternative molecular architectures in further depth. Inspired by the biological sequence of the NF-L protein, we synthesized sequence IDP^1^ to various hydrocarbon tails. By slightly modifying this sequence we studied how the interaction between the IDPs results in altered self-assembled structures once conjugated to the hydrophobic core.

IDPA^1^ series were synthesized with one or two aliphatic tails with three different tail lengths (1×14, 1×16, 1×18 and 2×12, 2×14, and 2×16) to investigate the influence of the hydrocarbon tail domain (Table 1, Supplementary Fig. S.1). In 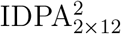, we segregated the negtaivly charged amino acids at the N-terminus, while the positive ones were placed at the C-terminus. Hence, both 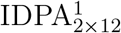 and 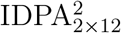 have identical magnitudes of net charge per residue (*NCPR* ≈ −0.278) at physiological pH.

Notably, the two peptide sequences include 11 chargeable residues, allowing for the net charge of the peptide to vary significantly as a function of pH. Electrostatic interactions are thus expected to play a significant role in the amphiphiles’ interactions and self-assembly. For both IDPAs, the isoelectric point (pI) is calculated at pH 4.1. At higher pHs, and in particular above pH 5.5, there is a decrease in the net charge to negative values due to the complete deprotonation of the aspartic acid and glutamic acid residues (Fig. S.15).

To investigate the role of single amino acid mutation, we designed 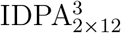, where we replaced the positively charged histidine at position 10 of 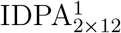 with neutral glycine (Supplementary Fig. S.1) which decreases the isoelectric point to 4.0. In previous experiments, we found that the hydrophilic domain (i.e., the disordered peptide) and its interactions controlled the complex aggregations at low pH and served to strengthen the interaction between worm-like micelles.^47^ Here, we focus on the intermediate pH region where a single mutation can potentially fine-tune the phase transition point.

The peptides’ degree of disorder was experimentally verified by measuring the circular dichroism (CD) spectrum (Supplementary Fig. S.3, see materials and methods). In addition, the free peptides, IDP^1^,IDP^2^ and IDP^3^ display a high probability for disorder and the absence of regular secondary structure using Iupred/Anchor^50^ and NetSurf 2.0^51^ algorithm (Supplementary Fig. S.4,S.5, S.6, for analysis methods see material and methods). Interestingly, changing a single amino acid (His to Gly at position 10) from IDP^1^ to IDP^3^ changes the pH-dependent disorder. Specifically, in the vicinity of the isoelectric point, IDP^3^ bioinformatiand CD analysis indicates a possible ordering and lack of disorder while IDP^1^ and IDP^2^ remain disordered throughout pH 2-10 (Supplementary Fig. S.4,S.5, S.6). Importantly, all the bioinformatic analysis is conducted on peptide sequences alone, assuming it is a good proxy for the IDPAs that contain hydrophobic domains. We verified this assumption by measuring the frequency resonance energy transfer (FRET) of Tyr at position 14 and Trp at position 1 of IDP^4^ and 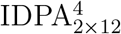. Here, we found no significant difference between the isolated peptide chain and when it is conjugated to the hydrophobic domain (Supplementary Fig. S.9, for FRET see material and methods).

### Amino-acids’ charge patterning regulates the self-assembled micellar structure at high pH

The self-assembly of each IDPA was characterized by measuring the structural properties of pH-equilibrated samples using an in-house and synchrotron small-angle X-ray scattering (SAXS). SAXS allows direct evaluation in the solution of both the nanoscopic self-assembled structures and the mesophase symmetry (Supplementary Information).

We began our self-assembly investigation by comparing the structures for 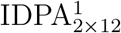 and 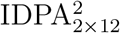 at pH 6.5, both having 2×12 hydrocarbon chains. In such conditions, both IDPAs self-assemble into a dispersed micellar state but with shifted SAXS patterns (Fig. 2). We fit the data using a spherical core-shell scattering form factor (Supplementary equ. (2)) and find that 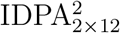 shows a significantly smaller radius 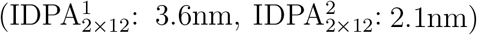. In addition, by extrapolating the form-factor to zero momentum transfer, we find the aggregation number to be about 40 and 20 for 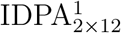 and 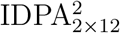, respectively. Given the similarity of the hydrophobic domain, the difference in radii thus originates from a smaller peptide layer (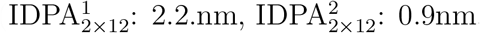, see Fig. 2a lower inset). Pair distance distribution function (PDDF) evaluation^55^ confirms that the 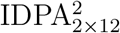 micellar phase has a significant smaller radius of gyration (Fig. 2a, upper inset). Notably, the assembled structure at pH 6.5 is robust with low polydispersity, indicative of the sharp SAXS features.

**Figure 2:**
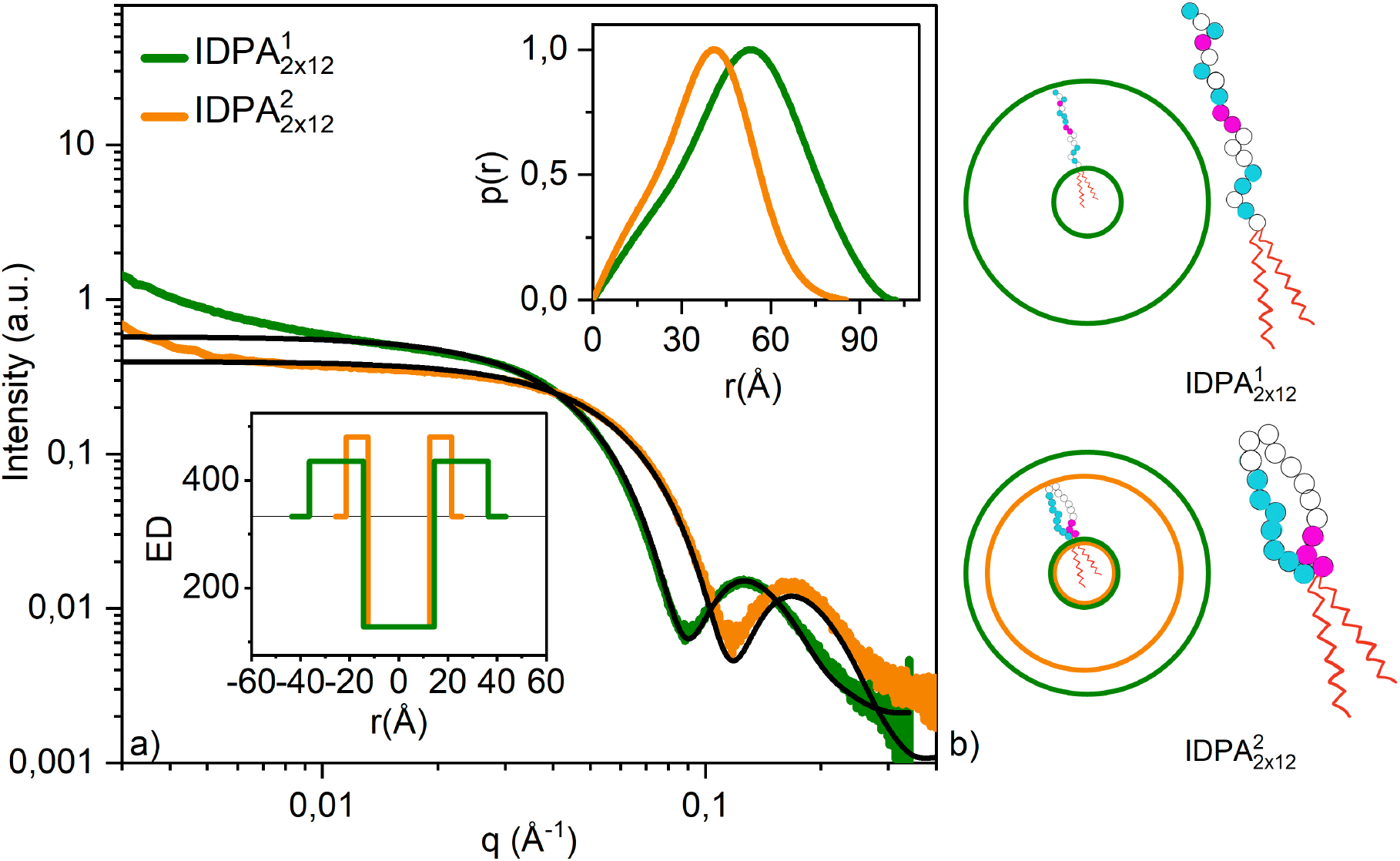
IDP head conformation as a function of sequence. a) SAXS profiles show smaller radii for 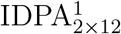 (green) than 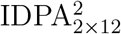 (orange) at pH = 6.35 ± 0.5. Micellar core-Shell form factor fits are shown in black line with parameters detailed in Supplementary table S.1. Lower inset: Electron density (ED) profile used in the fit. Upper inset: radius of gyration results of PDDF (*q*-range for fit: 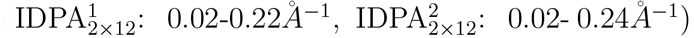. b) Representation of micellar sphere with 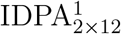 (green) and IDPA 2 (orange) with significantly different sizes of IDP layers and illustration of backfolding in 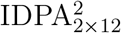 (lower cartoon) in comparison to 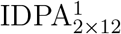 (upper cartoon). Pink circles indicate cationic, blue anionic, and white – neutral amino acids.

### Phase transitions are influenced by charged amino-acid positioning

Previous measurements^47^ of an amphiphile with a similar peptide head group as 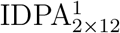 showed that its self-assembled structure is pH-dependent due to changes in the charged amino-acids. Here, we evaluate how charge patterning can tune the pH-dependent phase transitions for 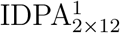 and 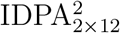 using SAXS, turbidity measurements, and cryogenic transmission electron microscopy (cryo-TEM). 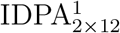 and 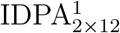 are insoluble close to the isoelectric point (pI). This indicates that peptide-peptide interactions are favored over peptide-water interactions.^56,57^ Away from the pI, the IDPAs become soluble and form monodisperse nanoparticles in the solution. These nanoparticles can be identified as spherical and/or cylindrical micelles using cryo-TEM and turbidity measurements (Fig. 3). Furthermore, SAXS data analysis and cryo-TEM direct imaging reveal that micellar rods collapse into a condensed phase in the vicinity of the pI (Fig. 3). For 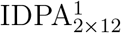, the SAXS data points towards a hexagonal phase (Fig. 3). This transition from worm-like monodisperse micelles to hexagonal packed was also studied by turbidity measurements showing a clear optical difference between the condensed and dispersed phase. Specifically, while 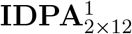 transitions in a relatively small pH interval (pH 4.2-4.6), 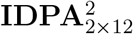 shows a significantly wider range for the transition (pH 4.2-6.5).

**Figure 3:**
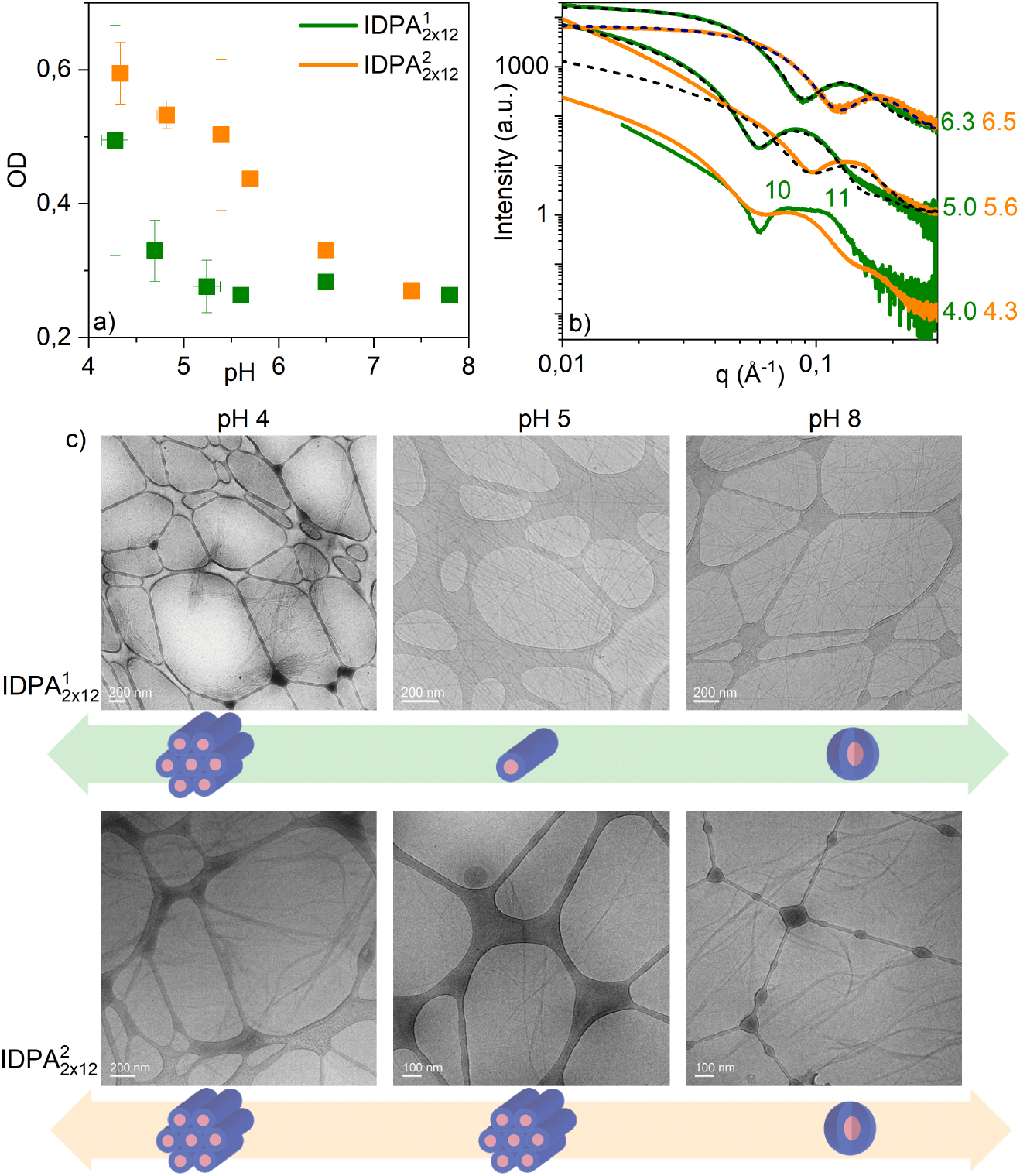
pH-dependent condensation of mesophases for 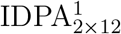 and 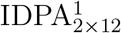 - from bulk to dispersed phase. a) Absorbance measurement shows high absorbance at the vicinity of the isoelectric point at pH 4.1, 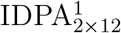 shows a significantly milder slope than 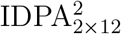 when transitions between the two states b) SAXS scattering for 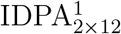 (green) and 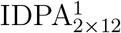 (orange) at various pHs. Dotted lines show spherical/worm-like core-shell form factors. 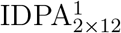 at pH 4 shows humps that point towards a hexagonal phase. CryoTEM pictures for 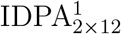 showing phase transition from spherical to worm-like micelles at pH 5. Aggregation of worm-like monodisperse micelles at the vicinity of the isoelectric point at pH 4.1.

### Both peptide sequence and hydrocarbon chains length tune the spherical to rod-like micelle transition

The balance between the architectures of the hydrophilic and hydrophobic domains plays a critical role in the self-assembly and phase transition of amphiphiles.^14^ Previously, we showed that hydrophobic dendritic domains conjugated to the peptide sequence of IDP^1^ could slightly alter the pH-induced phase transition from sphere to rod-like micelles.^47^ Here, we studied how the phase transition depends on the hydrocarbon length. Using SAXS, we find that double-chained 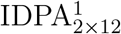 shows worm-like micelles at low pH and spherical micelles at high pH. At intermediate pH, we detect a coexistence regime with the combination of two mesophases by fitting the SAXS scattering through a linear combination of spherical and cylindrical core-shell shape factors (Fig. 4, and Fig. S.14). These results point to a continuous coexistence transition between spherical and worm-like micelles of constant radii. Significantly, the sharpness of the transition depends on the length of the tails: longer tails result in a phase transition at higher pHs with a much broader range (2×16: pH 4.7-7.8, 2×14: pH 4.7-7.5) between the two mesophases (Fig. 4). On the contrary, the 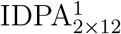 with shorter 2×12 tail transitions in a very narrow pH range (pH 5.7-6.0). Important to mention that for 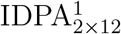 the cylinders transition completly to spheres whereas 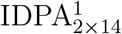 and 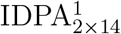 have still a low fraction of cylinders (approx. 2%) at high pHs.

**Figure 4:**
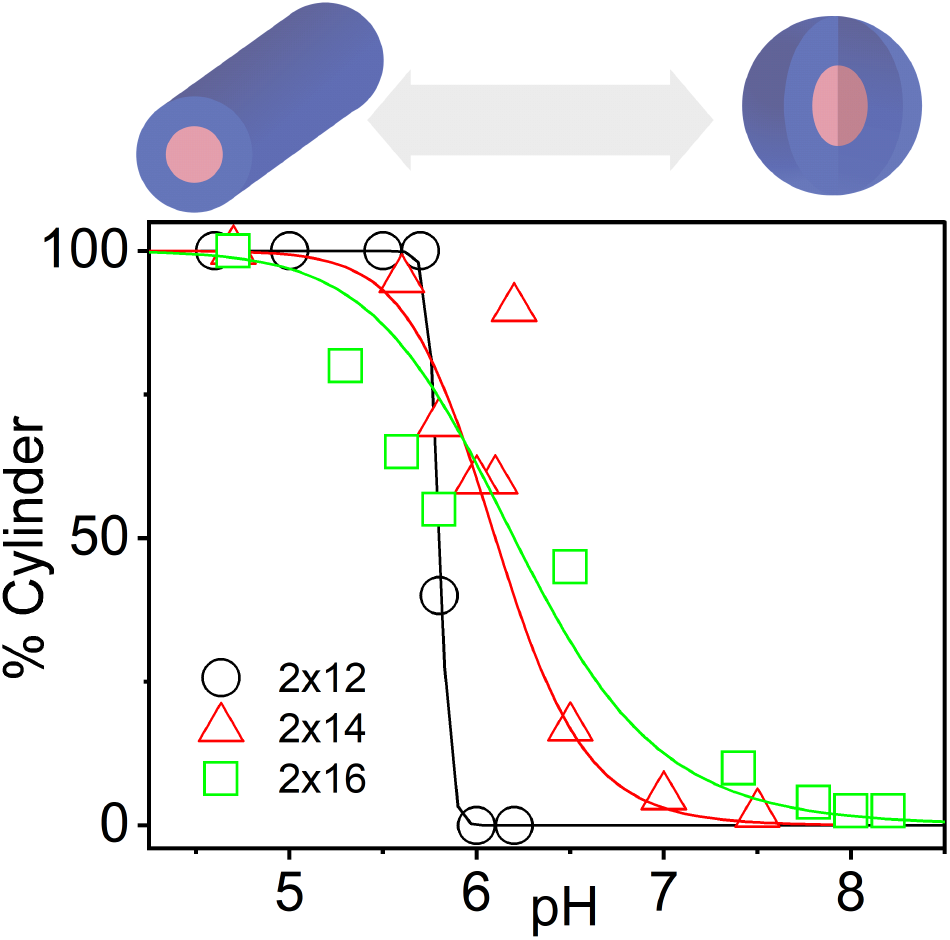
pH-dependent phase transition for different lipid chain length. The pH-dependent range of transition from worm-like (pH*<*4.7) to spherical micelles (pH*>*7.5) is broadening with increasing tail length (indicated in the legend). Phases in between are superposition of form factors and interpreted as coexistence. Lines represent a Hill function fit.

Using single value decomposition (SVD, see material and methods), we tested how many distinct scattering patterns contribute to the polydisperse signal for the transition pH range described before. We assume that the number of independent vectors resulting from the SVD analysis represents an upper bound to the number of different phases in the coexisting regime. Indeed, our IDPA transition requires up to 2-3 coexisting scattering vectors for the different IDPAs. Specifically, for IDPAs with tail lengths of 2×12 and 2×16, there are up to three different phases, and for the 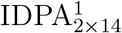, only two different phases are required by the SVD analysis (Fig. Sup. S.10). This result also agrees well with our initial finding that the IDPA transitions from spheres to rods, and in-between, we have a linear superposition of the two dominant form factors. For 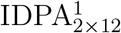 and 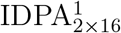, we found that three independent vectors can describe the data. A possible explanation is an intermediate phase, e.g., an ellipsoidal phase, between the rod and the spherical phase that, unfortunately, is too weak for us to fit even by synchrotron’s SAXS data.

The number of hydrocarbon chains is another architectural feature when designing IDPAs. For double-chained IDPAs, the SAXS pattern is isotropic as the nanoparticles scatter of all possible orientations (Fig. 5a). However, while IDPAs with single hydrocarbon tails (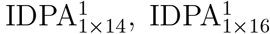 and 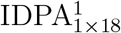) show isotropic micellar spheres at intermediate and high pHs, they collapse into liquid crystals with a strong “spackle” pattern close to the pI. The scattering peak positions indicate Face Centered Cubic (FCC) and Body-Centered Cubic (BCC) Bravais lattices (Fig. 5b). Importantly, around the pI, the FCC and BCC organizations and “spackle” scatterings are evidence of soft IDPA monodispersed micelles packed into rather large “crystals” on the incoming beam dimensions (≈ 1.5 mm^2^). The SAXS analysis reveals that the lattice parameters for both FCC and BCC are proportional to hydrocarbon tail lengths (𝓁, see Fig. 5c and d). Using 𝓁 *<* 𝓁 _*max*_ = (1.54 + 0.1265*nm*) as an approximation for hydrocarbon tail extension,^17^ we can extract the approximate size of the hydrophilic domain thickness to be around 2.7*nm*. The hydrophilic domain thickness does not depend on the hydrocarbon tail length. Moreover, the IDP layer at the isoelectric point is in agreement with the IDP layer of micellar spheres fitted at intermediate pH (Supplementary table S.1 and Fig. S.13 and dashed lines in Figs. 5 c,d).

**Figure 5:**
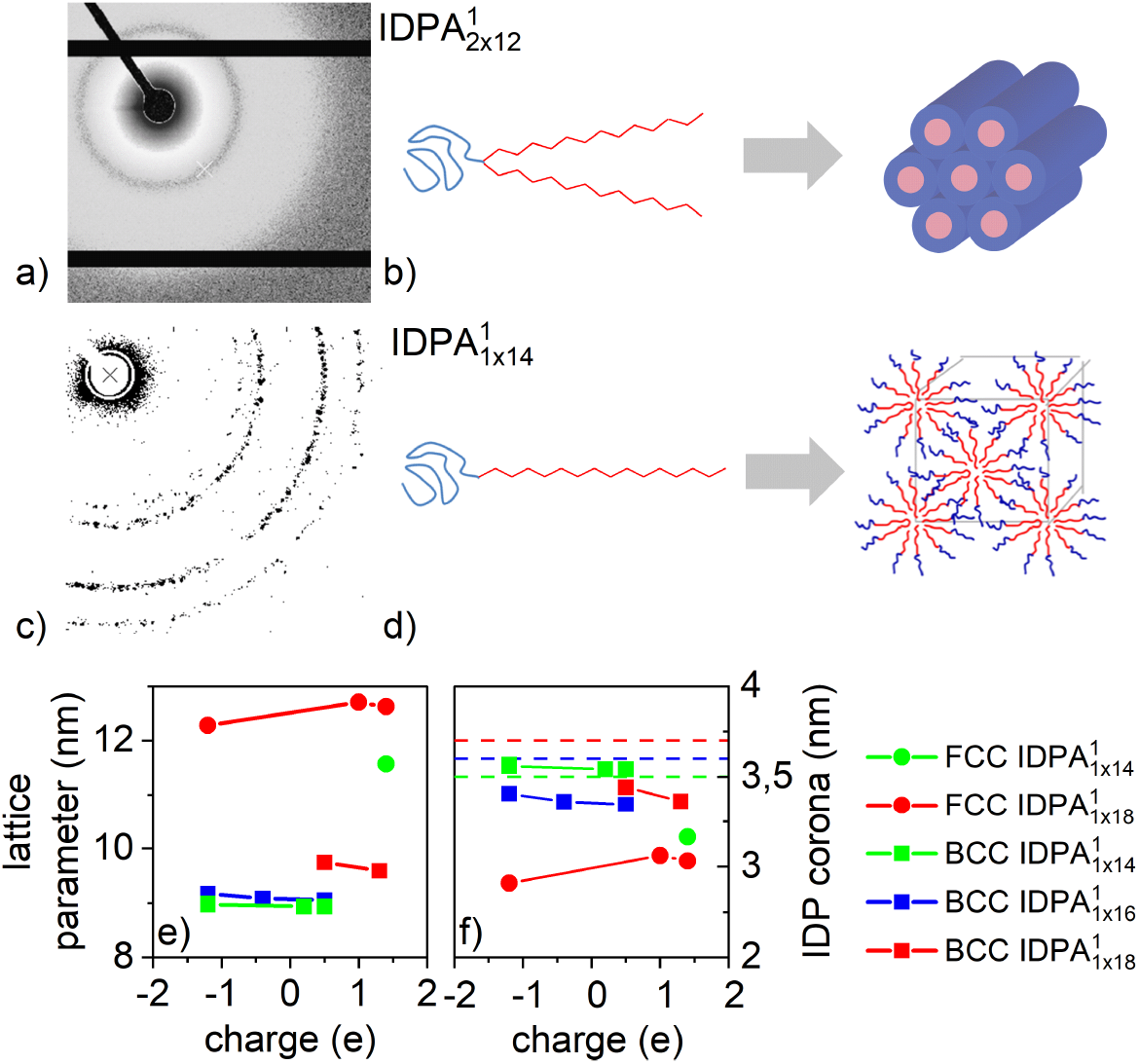
Formation of liquid crystals at isoelectric point (pH 4.3) for single tailed IDPAs with different tail lengths. 2D SAXS pattern for (a) double and single (c) tailed IDPAs at isoelectric point (pH 4) showing hexagonal and FCC phases. (b) and (d) are related cartoons illustrating the formation of mesophases from the double and single tailed IDPAs, respectively. Lattice parameters (*d*) for (e) BCC and (f) FCC phases from integrated 1D patterns for single tailed IDPAs near isoelectric point were found by extracting peak position via gaussian fit. The charge is calculated via summation of amino acids’ charges at various pHs. Unit cell dimensions are directly-measured from SAXS correlation peak positions. Nearest neighbours (dashed lines) are extracted using 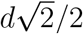 for FCC and *d* for BCC. IDP headgroup layer sizes for 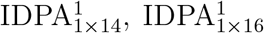 and 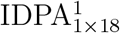 are extracted by subtracting the calculated tail length (𝓁 _*max*_, see text) from the lattice parameter.

After studying how the length and the number of tails affect the self-assembly of the IDPAs, we set to explore how minor alterations in the peptide sequence can tune the phase transition. For example, 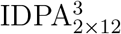, which is different from IPDA1 only by single amino acid at position 10, transitions at pH 5.4 from spherical to cylindrical micelles, while the equivalent 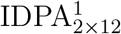 transitions at pH 5.8 (Supplementary Fig. S.15). The altered transition can be attributed to differences in interactions resulting from exchanging Histidine (*pK*_*a*_ = 6.0) with the neutral glycine. An alternative route to influence the self-assembly is through the introduction of salt (NaCl), which screen the electrostatic interactions between neighboring charged peptides. Using Kratky analysis on the SAXS data, we reveal the compactness of the IDPAs at varying salt concentrations (Supplementary Fig. S.11). We find a trend towards higher slopes in the high momentum vector (*q*) region with increasing salt concentration. This is more pronounced with increasing chain length. The high slope indicates that the IDPAs are more unfolded than at low salt concentrations. For the low *q*-region, the dispersity between the curves becomes more pronounced with increasing chain length.

### Cleavable IDPAs

One of the advantages of IDPAs is the ability to design sequences that can interact with other biological entities. For example, the utilization of IDP as the hydrophilic domain can be designed to interact with an enzyme, in order to induce drug release from the self-assembled nano-carrier or aggregation of the carrier at the site of enzymatic activation.^58,59^ Therefore, we designed the additional IDPA sequence (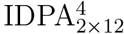, Supplementary Fig. S.1) that contains a cleavage domain (GPLGLAG) for an MMP-9 enzyme. Indeed, upon incubation with the MMP-9 enzyme, the IDPA is cleaved with a shortened peptide sequence (Supplementary Fig. S.2). We term the remaining amphiphile, which includes the hydrophobic domain, as 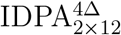 and the cleaved peptide as IDP^4*δ*^.

The cleavage site in 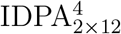 was introduced to disturb the self-assembled structure via enzymatic reaction dramatically. The sequence conjugated to the hydrocarbon (IDP^4Δ^) contains neutral amino acids. It is on the threshold of being disordered, while the remaining part (after the cleavage site), termed here IDP^4*δ*^, contains partially protonatable amino acids and is expected to be disordered at all pHs (Supplementary Fig. S.7, S.8). Both, 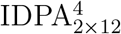 and 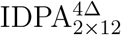, were measured at various pHs, and their self-assembly was studied using SAXS.

At physiological pH, 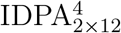assembles into spherical micelles, indicated through the scattering intensity at small angels,^60^ *I*(*q* → 0) ∼ *q*^*−*0^, while 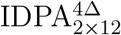is forming worm-like micelles with *I*(*q* → 0) ∼ *q*^*−*1^ (Fig 6 a). We further fit the SAXS data using a (smooth) spherical core-shell model and a cylindrical core-shell model and found that the hydrocarbon domain stays constant while the peptide layer of the sphere is smearing toward higher radii with lower electron densities (Fig 6b).

**Figure 6:**
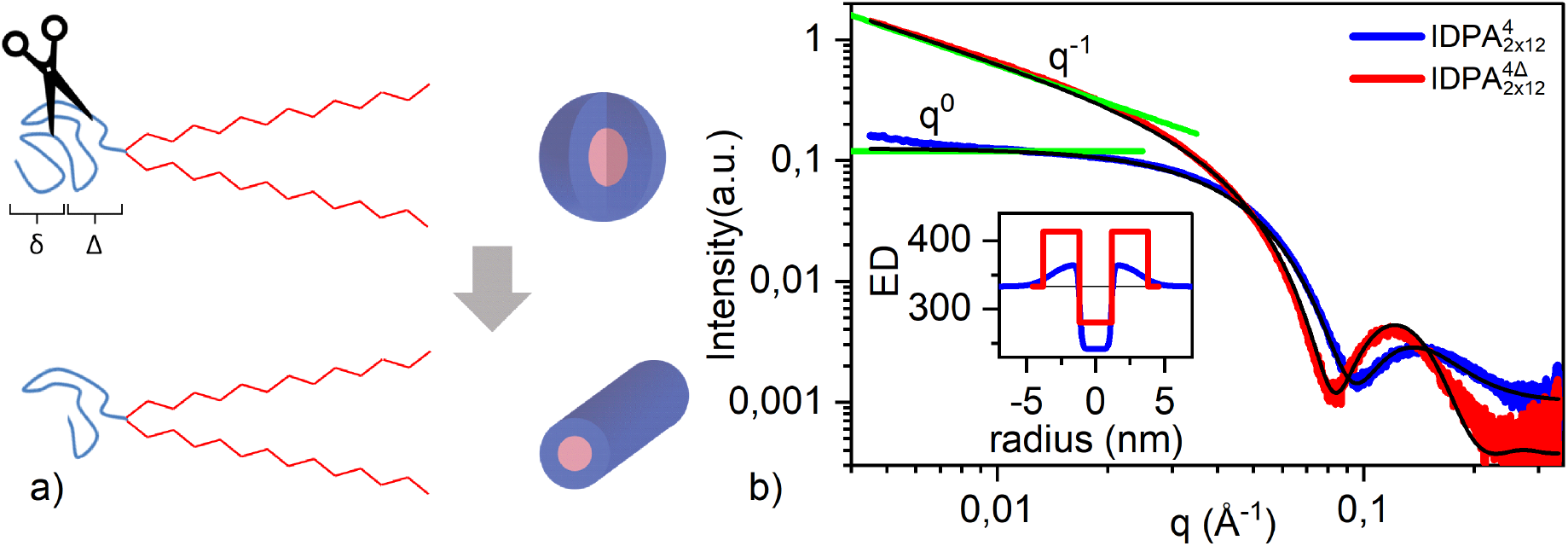
SAXS data for cleavable IDPA. a) Cartoon showing self-assembly of spherical and worm-like micelles for 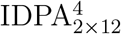 and 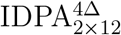, respectively. b) SAXS data and fit for the IDPAs (blue 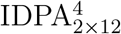, red 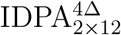) at physiological pH (pH 7). Inset shows electron density (ED) profiles. Green lines show small angle region fits used for initial structural determination.

At different pH values, 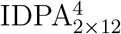 undergoes structural phase transitions while 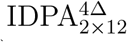 remains in a worm-like state (Supplementary Fig. S.16). In agreement with Takahashi et al.,^61^ the SAXS pattern for 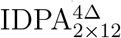 at pH 5 indicates the formation of polymer vesicles upon stretching of spherical micelles.

Furthermore, this pH sensitivity was investigated by measuring the *R*_*g*_ versus the pH of the crude peptides using SAXS. The *R*_*g*_s for IDP^4^ and IDP^4Δ^ show little dependence on pH and are ∼ 9 and ∼ 11*Å*, respectively. However, we assume that the *R*_*g*_ of IDP^4^ is more sensitive to pH (Fig. S.17). This is indicative of the pH-sensitive phase change of 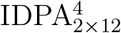 compared to 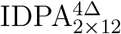.

## Discussion

IDPAs present a highly modular molecular platform for the design of transformative nanocarriers.^47^ We presented new IDPA molecules, which were entirely synthesized by an automated solid phase peptide synthesizer. A peptide sequence inspired by the disordered regions of neurofilament light chain protein was systematically altered to study how the interplay of hydrophobic tail(s) architecture and polypeptide headgroup conformation dictates the self-assembly process.

Despite sharing identical amino acids, 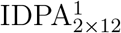 and 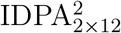, with similar hydrophobic domains, assemble into spherical micelles with different radii at high pH. Specifically, 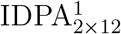 assembly has a significantly larger polypeptide shell thickness than 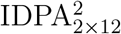.

This demonstrates how to sequence ordering plays a dominant role in the assembly of ID-PAs. In 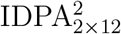, we segregated the positive and negative charged amino acids at the edges of the sequence. Therefore, the more compact peptide conformation is likely to result from transient back folding of the peptide chains due to electrostatic interactions of the oppositely charged ends (Fig. 2b).

Investigation of the self-assembly of the two IDPAs at different pH values revealed that the transition from a collapsed hexagonal phase at the isoelectric point to dispersed worm-like micelles is also sequence-dependent. For example, 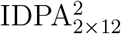 transitions to a dispersed state over a relatively broad pH range compared to 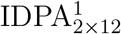. Considering our previous results,^47^ we argue that the transient hairpin-shaped and more compact peptide conformation are less-prone to interact with neighboring worm-like micelles. In a sense, for 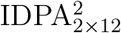, the almost complete overlap between the peptides of opposing worm-like micelles is needed to induce electrostatic attraction, while for 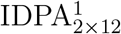, only partial overlap is needed.

In addition, even a minor alteration, such as the exchange of a single amino acid in the peptide sequence, can tune the pH structural phase transition. Specifically, 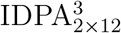 transitions between spheres to elongated worm-like micelles at pH 5.4, while 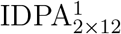 transitions at pH 5.8 with a change of one single amino acid (histidine to glycine). When calculating the net charge difference between 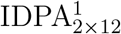 and 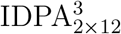, one can expect that the phase transition will occur at pH 5.2 (Supplementary Fig. S.15a), although experimentally, the difference is milder. Using a free energy model for electrostatic repulsion contribution, we can explain this phenomenon.^47^ In short, the position of the charged amino acid along the polypeptide contributes to the electrostatic repulsion between the neighboring chains in proportion to their vicinity to the peptide-tail interface. Therefore, exchanging the charged histidine in the middle of the sequence has a relatively mild impact on the mesoscopic structural phase transition.

As an alternative means to alter the structural phase transition, we evaluated the role of the hydrophobic tail(s) domain. When introducing IDPAs with just one chain instead of two, the IDPAs self-assembled into large spherical micelles crystals close to the isoelectric point. As shown in Fig. 5, the distance between these micelles within the crystals is significantly smaller than the micelles radii at slightly higher pHs. This indicates that the outer IDPs’ shells overlap between nearest neighbors. Such overlap is needed to induce short-ranged attractive forces between neighboring IDPAs, stabilizing the micellar crystals.

At intermediate pHs, the IDPAs are in the coexistence phase of spheres and cylinders, where the transition width broadens with increasing tail length. While similar coexistence of rod and micellar phases, instead of elongated micelles with end caps, has been shown before,^62^ the correlation between the transition width and the chain length requires further explanation, as detailed below.

It was proposed that the reason for the coexistence between cylindrical micelles of finite lengths and spherical micelles is an energy barrier the system has to overcome on the way of transformation between the two types of micelle.^62^ This energy barrier originates from the difference between the energy of two endcaps of a cylindrical micelle and the energy of a spherical micelle. Hence, such coexistence does not represent a thermodynamic equilibrium between the two phases but rather indicates a slow transition between the two phases enabling simultaneous observation of both cylindrical and spherical micelles within the time scale of the experiments. In this model, the beginning of the coexistence region (Fig. 4) corresponds to conditions upon which the energy barrier of formation of a spherical micelle out of a cylindrical one is such that the characteristic time of this event is comparable with the time of observation. At the end of the coexistence region, the energy barrier must be small enough to make the transition time shorter than the observation time. The origin of the energy barrier is an energetically unfavorable but unavoidable transition region, which builds up within a cylindrical micelle between its endcap and the cylindrical part because of a difference in their cross-sectional thicknesses.^62^ This difference results from packing molecules with a particular molecular volume and surface area into a spherical versus cylindrical aggregate. An increase in the spontaneous monolayer curvature driven by the charge growth at increasing pH makes the endcap more energetically favorable and hence decreases the energy barrier. A simple geometrical consideration explains that the shorter the IDPA chain length, the more minor the thickness mismatch between the endcap and the cylindrical part of a micelle and, therefore, the lower the initial energy barrier. As a result, less charge must be generated to cut down this energy barrier and facilitate a fast cylinder-to-sphere transition. This explains the chain length dependence of the width of the coexistence region (Fig. 4).

We have shown that IDPAs can be engineered to induce phase transitions upon enzymatic activation. 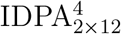 self-assembles into spherical micelles, whereas upon enzymatic cleavage, the assembly of the cleaved 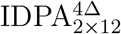 transforms into worm-like micelles at physiological pH. Furthermore, we demonstrated that pH triggers phase transitions for the uncleaved peptide containing protonable amino acids, whereas pH does not affect the cleaved peptide containing only neutral amino acids. These results are of great interest for biomedical applications, given the ability to change the physical properties of the nanocarrier at constant pH by an enzymatic reaction. It thus suggests an alternative path for enzymatically triggered activation of drug release in a controllable manner. Furthermore, 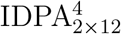, in similarity to all other IDPAs presented here, shows remarkably controllable, monodisperse nano-structures. The pH dependency of 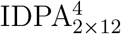 and 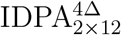 self-assembly demonstrates the ability to design both pH-dependent and independent structures upon cleavage. Thus, our work enables us to combine enzymatic cleavage with pH-dependent phase transition in a single amphiphilic molecule.

## Conclusion

We have studied the self-assembly of five disordered polypeptide domains conjugated with different fatty acids in IDPAs. Even though polypeptide chain conformation is disordered, the interactions between the peptide headgroups lead to various distinct self-assembled nanostructures. The IDPA systems respond to pH and salinity and exhibit structural phase transitions depending on the peptide sequence and the number and length of the hydrocarbon tail.

It stands to reason that IDPAs mesostructures such as micelles, micellar tubes, or condensed phases and their defined structural transitions could potentially be exploited in biotechnological applications or as drug delivery nanomaterial in biological environments.

In this context, it is notable that pH-dependent phase transitions are sensitive to single amino acid mutations within the sequence. The width of the structural phase transition can be tuned by choosing hydrocarbon tails.

Furthermore, it is remarkable that permutations in the amino acid sequence led to different average conformations, e.g., extended or transient hairpin-like back folding. Thus, disordered peptide motifs can result in distinctly different average conformations dependent on amino acid composition and sequence order. Last, we designed an enzymatically cleavable IDPA to demonstrate that IDPAs as surface-active components of nanocarriers can potentially react to metabolic conditions at target sites.

IDP based headgroups may serve as grafted polymers for stabilizing particles via shell-formation, as alternative to polyethylene glycol (PEG)-lipids. Overall, their highly modular structure and function make IDPAs valuble to implement tailored functionalities and finetuned interactions for controllable structural phase transitions that could expedite cargo release. Based on our results and the discussed advantageous properties, we expect that IDPA conjugates will be valuable resources for the research community advancing precision medicine.

## Supporting Information

Chemical formulas, disorder analysis, FRET data, SAXS data and details about SAXS analysis

## Acknowledgement

The synchrotron SAXS data was collected at beamline P12 operated by EMBL Hamburg at the PETRA III storage ring (DESY, Hamburg, Germany), and the SOLEIL synchrotron facility for time on Beamline SWING and at beamline B21 at Diamond Light Source. We would like to thank Haydyn Mertens (DESY), Thomas Bizien (Soleil), Nathan Cowieson and Charlotte Edwards-Gayle (Diamond Light Source) for the assistance in using the beamlines. This work was supported by the National Science Foundation under Grant No. 1715627, the United States-Israel Bi-national Science Foundation under Grant No. 2020787, and the Israel Science Foundation under Grants No. 1454/20. We also acknowledge fruitful discussions with Vladimir Uversky, Ram Avinery and Uri Raviv.

## Supporting Information

This file contains:

- Chemical structures (Fig. S1) and tables with key parameters for fitting
- Figures S2-S12 referred in the main text
- Form factor equations for SAXS, including figures Sx referred here

