## Supplementary information for "Self-Assembly of Tunable Intrinsically Disordered Peptide Amphiphiles"

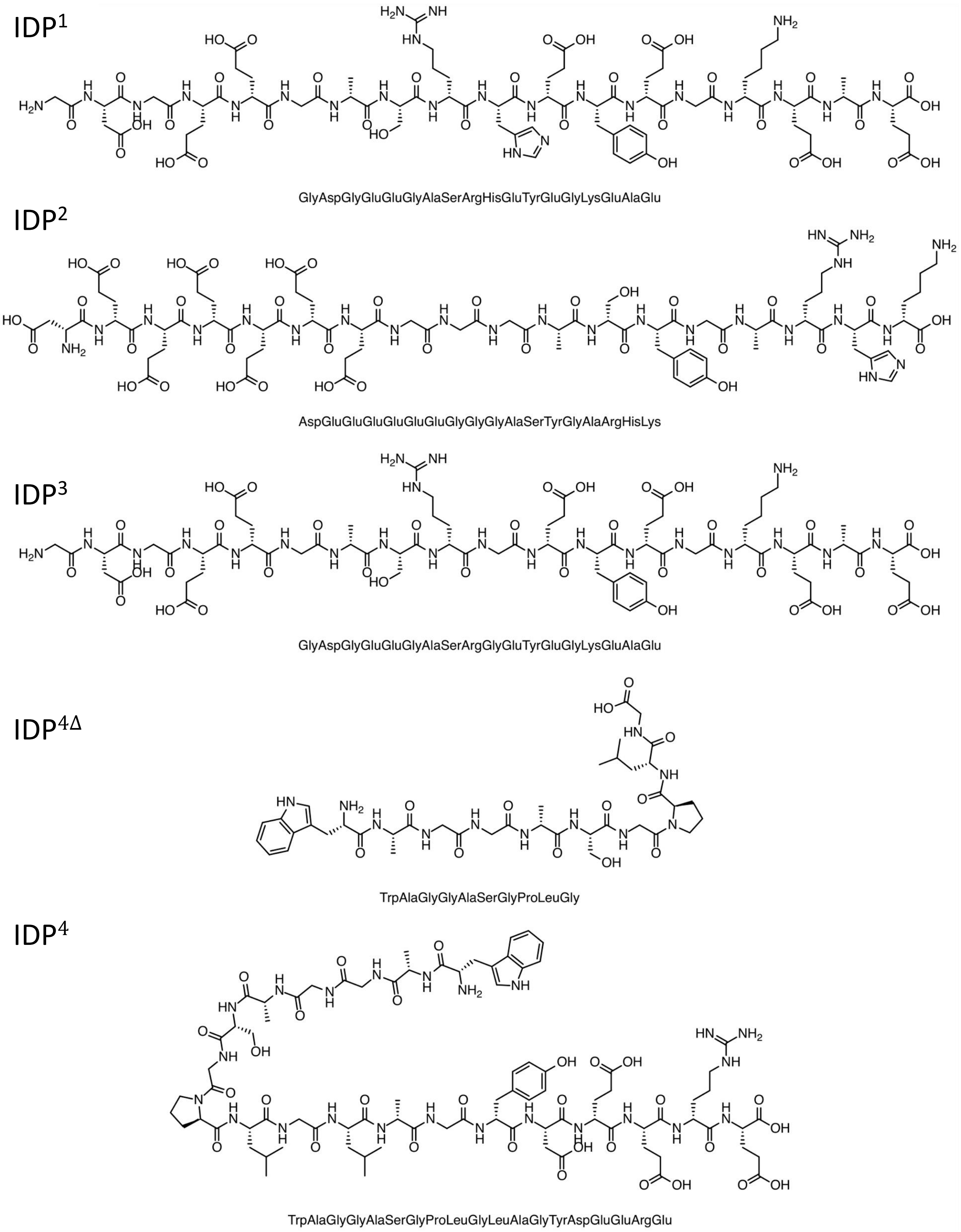


Figure S.1: **Molecular structure** for peptide sequences shown for IDP$^1$,IDP$^2$, IDP$^3$, IDP$^{4\Delta}$ and IDP$^4$, below three letter code

Table S.1: **Fitting parameters**. Radially integrated SAXS data was fitted with either a spherical or cylindrical core shell model using X+ (Supplementary information, and[63]). $r_{total}(\AA)$ describes the radii of the total spherical/worm-like micelles while $r_{core}(\AA)$ and $r_{shell}(\AA)$ show the value of the radii of the hydrocarbon core and the peptide layer, respectively. $SLD(enm^{-3})$ stands for scattering length density and describe the amount of scattering electron per certain volume. While $r_{core}$ stays rather constant, worm-like micelles have a significant bigger $r_{shell}$

| | Spherical micelles (high pH) | | | | | Worm-like micelles (low pH) | | | | |
|---|---|---|---|---|---|---|---|---|---|---|
| | $r_{total}$ ($\AA$) | $r_{core}$ ($\AA$) | $r_{shell}$ ($\AA$) | $SLD_{core}$ ($enm^{-3}$) | $SLD_{shell}$ ($enm^{-3}$) | $r_{total}$ ($\AA$) | $r_{core}$ ($\AA$) | $r_{shell}$ ($\AA$) | $SLD_{core}$ ($enm^{-3}$) | $SLD_{shell}$ ($enm^{-3}$) |
| $IDPA^{1}_{2\times12}$ | 36±1 | 14±1 | 22.0 | 139±6 | 418±20 | 54±1 | 14±1 | 40±1 | 271±5 | 395 ±20 |
| $IDPA^{1}_{2\times14}$ | 36±1 | 14±1 | 22,50 | 135±7 | 413±23 | 56±1 | 16±1 | 40 ±1 | 271±5 | 395 ±20 |
| $IDPA^{1}_{2\times16}$ | 36±1 | 25±1 | 21±1 | 158±30 | 428±8 | 57±1 | 17±1 | 40 ±1 | 271±5 | 395 ±20 |
| $IDPA^{1}_{1\times14}$ | 38±1 | 13±1 | 25±1 | 228 ±23 | 392 ±17 | | | | | |
| $IDPA^{1}_{1\times16}$ | 40±1 | 14±1 | 26±1 | 218 ±21 | 408 ±12 | | | | | |
| $IDPA^{1}_{1\times18}$ | 46±1 | 18±1 | 27±1 | 218 ±21 | 403 ±19 | | | | | |
| $IDPA^{2}_{2\times12}$ | 21±1 | 13±1 | 9±1 | 481±6 | 128±21 | | | | | |
| $IDPA^{4}_{2\times12}$ | | | | | | 36±1 | 12±1 | 26±1 | 280±5 | 414±16 |
| $IDPA^{4\Delta}_{2\times12}$ | 33±1 | 12±1 | 21±1 | 242±15 | 367±1 | | | | | |

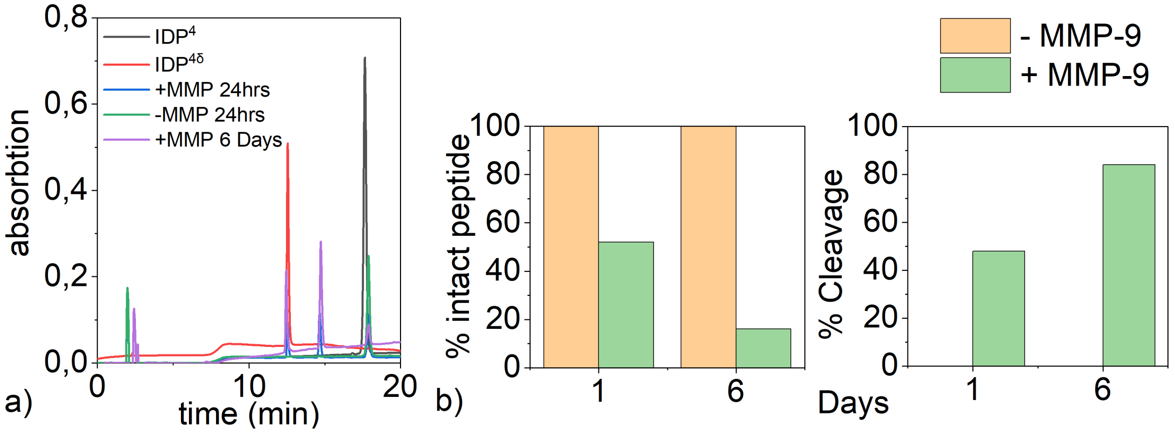


Figure S.2: **Enzymatic cleavage with MMP-9 Enzyme** a) HPLC was run on $IDP^4$ and $IDP^{4\delta}$ to determine their retention times. (Experiment Details: Buffer: 200 mM NaCl, 50 mM TRIS, 5 mM CaCl2, 1 mM ZnCl2, pH = 7.5, peptide conc. = 85$\mu$M, enzyme conc. = 0.85$\mu$M, total Volume = 100 $\mu$l, MMP-9 Enzyme: lot # 09051736)). Peptide solution was treated with MMP-9 enzyme and HPLC was obtained at 24 hrs and 6 days. Control samples without addition of MMP-9 were also run to check for non-specific cleavage over the reaction time. b) Diagram for intact and cleaved peptides after 1 and 6 days.

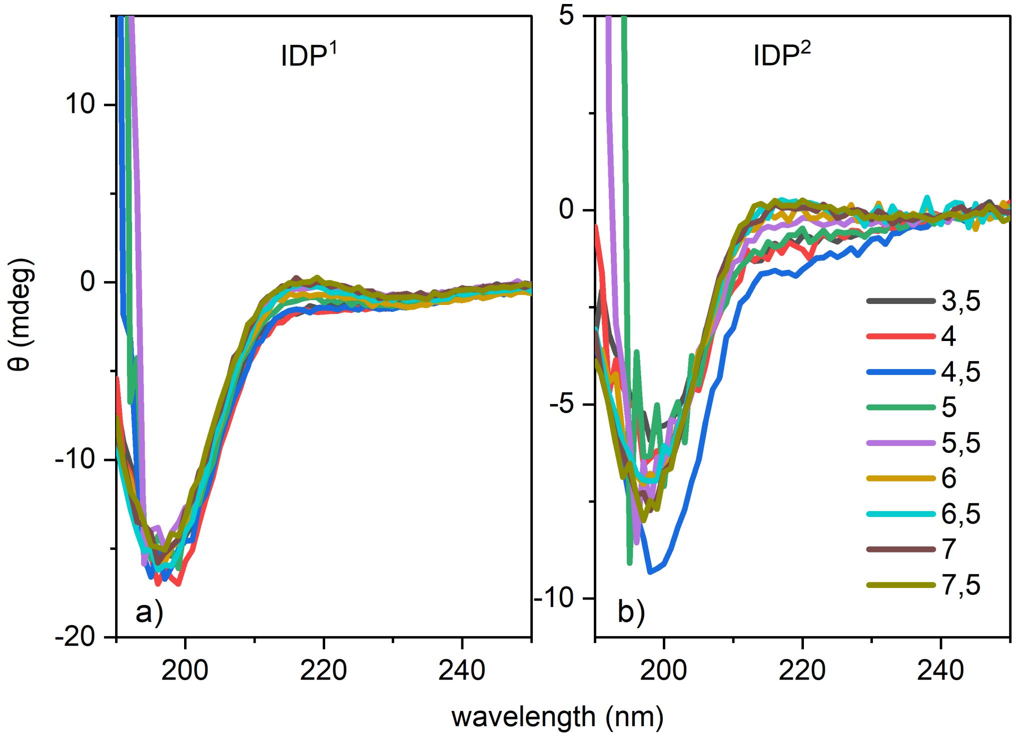


Figure S.3: **CD measurements** of IDP[1] and IDP[2] in phosphate buffer (the buffer was changed due to strong absorption of the normal buffers used for sample preparation). Peptide was measured in the presence of 10mM sodium acetate, and sodium phosphate buffer for pH 3.5-7.5, respectively. CD signal present random coil spectrum for all IDPs which indicates unstructured peptides (disordered) and no secondary structure for the relevant pHs.

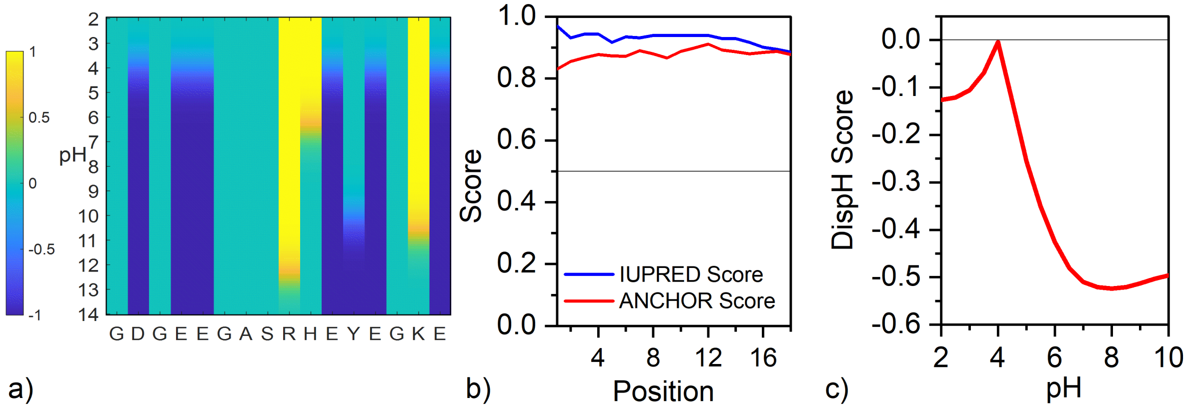


Figure S.4: **Charge and disorder analysis for IDP[1]** a) Charge state for each amino acid at different pHs. b) Iupred/Anchor analysis shows disorder tendency (threshold: 0.5) for IDP,[50] further analyis with Netsurf supports this statement.[51] (c) pH dependent disorder analysis with DispHred[64] predicts disorder tendency for all pHs.

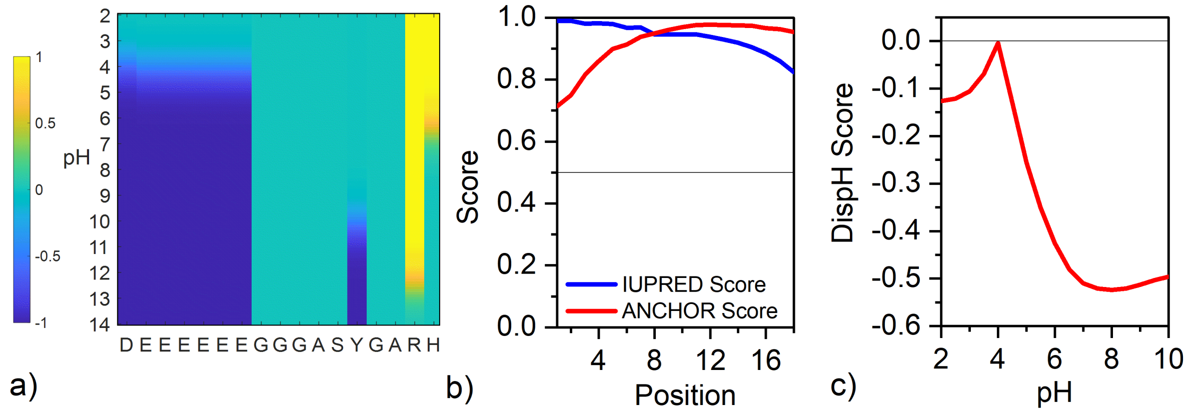


Figure S.5: **Charge and disorder analysis for IDP**[2] a) Charge state for each amino acid at different pHs. b) Iupred/Anchor analysis shows disorder tendency (threshold: 0.5) for IDP,[50] further analyis with Netsurf supports this statement.[51] (c) pH dependent disorder analysis with DispHred[64] predicts disorder tendency for all pHs.

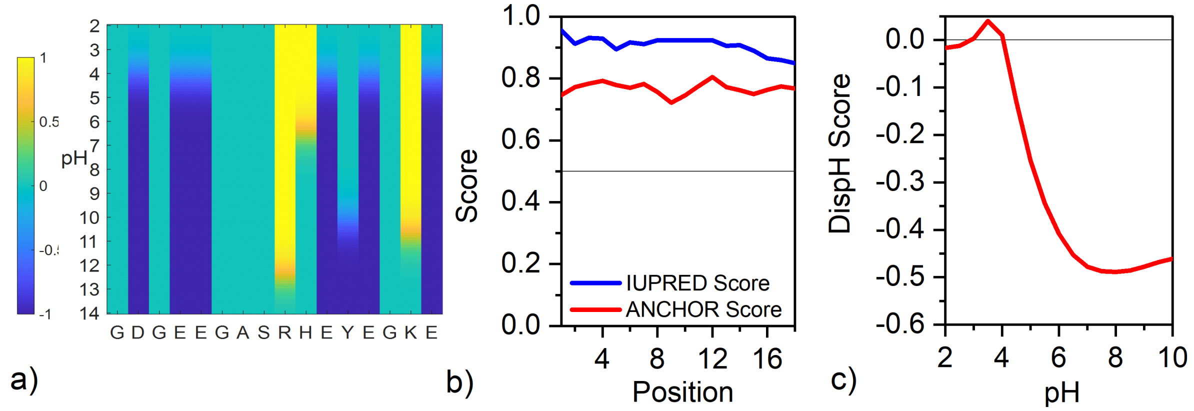


Figure S.6: **Charge and disorder analysis for IDP**[3] a) Charge state for each amino acid at different pHs. b) Iupred/Anchor analysis shows disorder tendency (threshold: 0.5) for IDP,[50] further analyis with Netsurf supports this statement.[51] (c) pH dependent disorder analysis with DispHred[64] indicates a possible ordering and lack of disorder the vicinity of the isoelectric point.

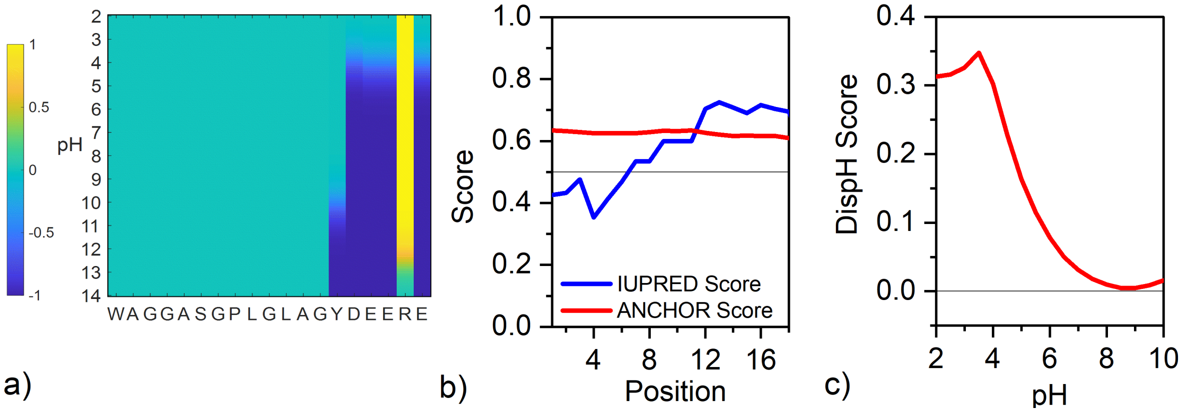


Figure S.7: **Charge and disorder analysis for IDP[4]** a) Charge state for each amino acid at different pHs. b) Iupred/Anchor analysis shows disorder tendency (threshold: 0.5) for amino acids after the cleavage site,[50] further analysis with Netsurf supports this statement.[51] (c) pH dependent disorder analysis with DispHred[64] predicts disorder tendency for all pHs.

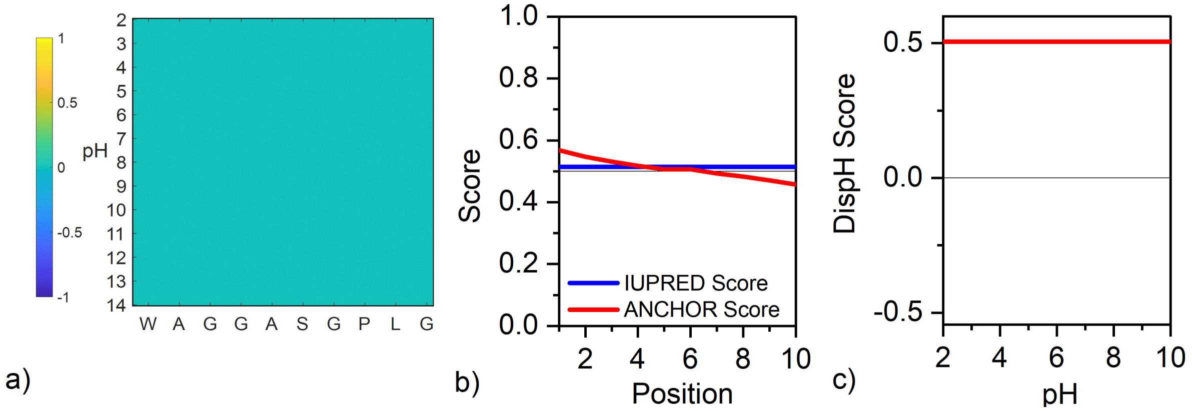


Figure S.8: **Charge and disorder analysis for IDP$^{4\Delta}$** a) Charge state for each amino acid at different pHs. b) Iupred/Anchor analysis shows that IDP is on the threshold to be disordered,[50] further analysis with Netsurf supports this statement.[51] (c) pH dependent disorder analysis with DispHred[64] predicts no disorder tendency for all pHs.

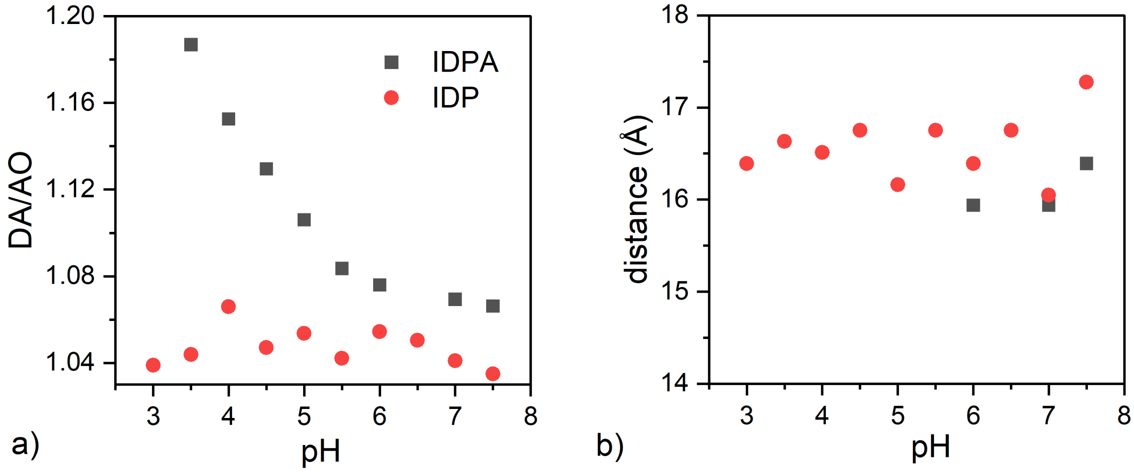


Figure S.9: **FRET measurements of IDP$^4$ and IDPA$^4$** a) Fluorescence ratios for IDP$^4$ and IDPA$^4$ b) Extracted distances show that the distance between Tyr at position 14 and Trp at position 1 is not changing through the pH range of 3.5 – 7.5. A small decrease in the distance was found in IDPA$^4$ compared to IDP$^4$ at pH 6.5-7.5. We were not able to calculate the energy transfer at lower pH of IDPA$^4$ due to change in the spectrum profile of the WY dipeptide reference. But, an increase in the fluorescence spectra for IDPA4 was seen in the lower pH 3.5-6.5 suggesting a decrease in the donor-acceptor mean distance.

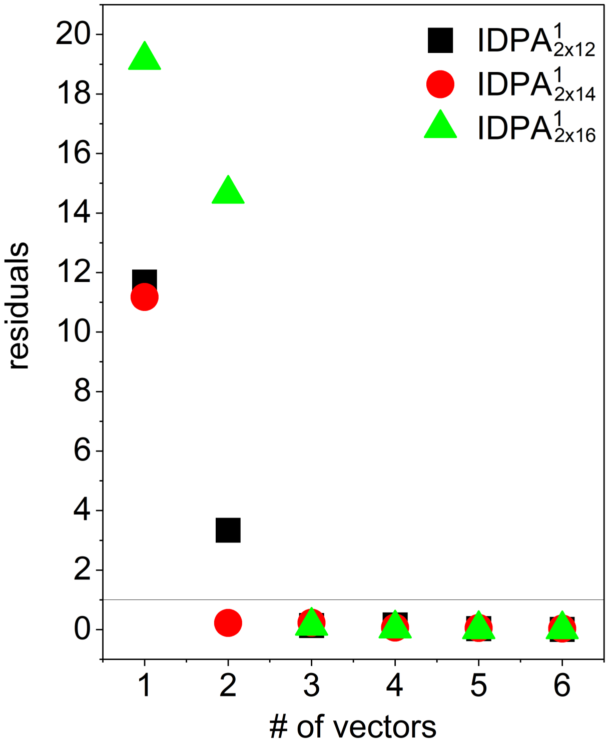


Figure S.10: **Singular value decomposition (SVD) analysis** of pH dependent SAXS data for spherical to worm-like micelle transition. Analysis followed the SVD analysis. The residuals $R_k = \frac{1}{nm}\sum_{q_i=1}^{n}\sum_{p_j=1}^{m} R^2_{k,q_i,p_j}$, where $m$ is the size of the scattering vector $q$ and $n$ are the number of pH steps, are plotted as a function of the number of singular vector components $(k)$ that were chosen to reconstruct the data matrix. Here, $R_{k,q_i,p_j}$ was defined by $R_{k,q_i,p_j} = \frac{D_{q_i,p_j} - D_{kq_i,p_j}}{\sigma_{q:i,p_j}}$, where $D$ is the data matrix, in which each column represents a one dimensional scattering curve, $I(q,p)$ at every pH step $p$. $D_k$ is the reconstructed data matrix using $k$ singular orthonormal vectors, and each term $(q_i, p_j)$ in the matrix $\sigma$ corresponds to the measured standard error for the corresponding term in $D$. The black line indicates the cutoff where the residual is equal to 1.[65]

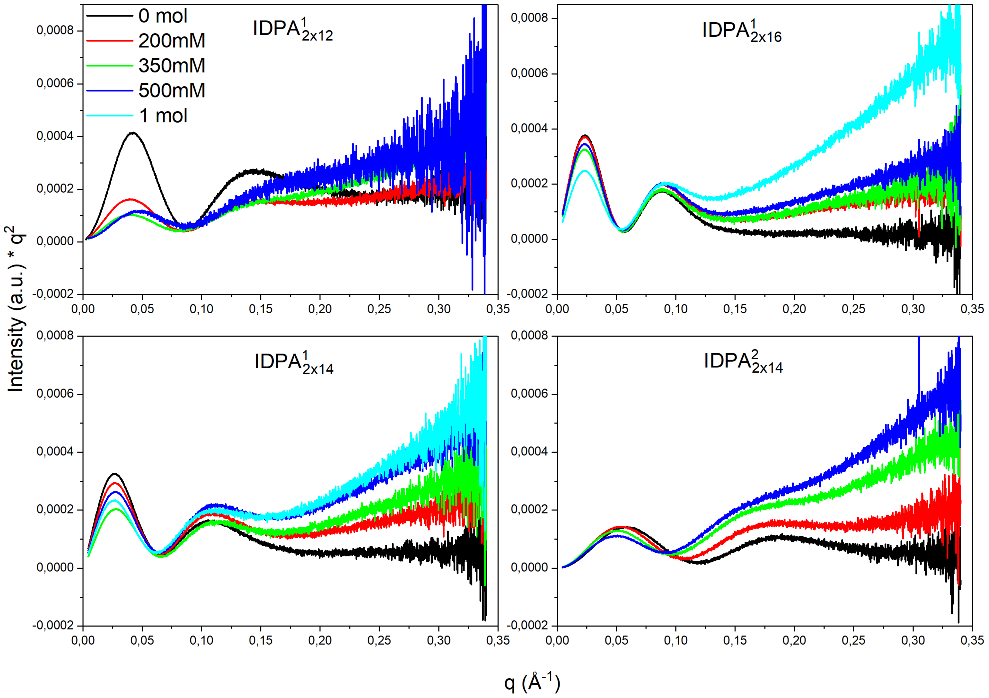


Figure S.11: **Kratky Plots for IDPAs with different salt concentration**. Scattering in high angles become more pronounced with increasing salt. Higher chain length strengthens the trend. For low q, polydispersity between plots becomes more pronounced with increasing chain lengths.

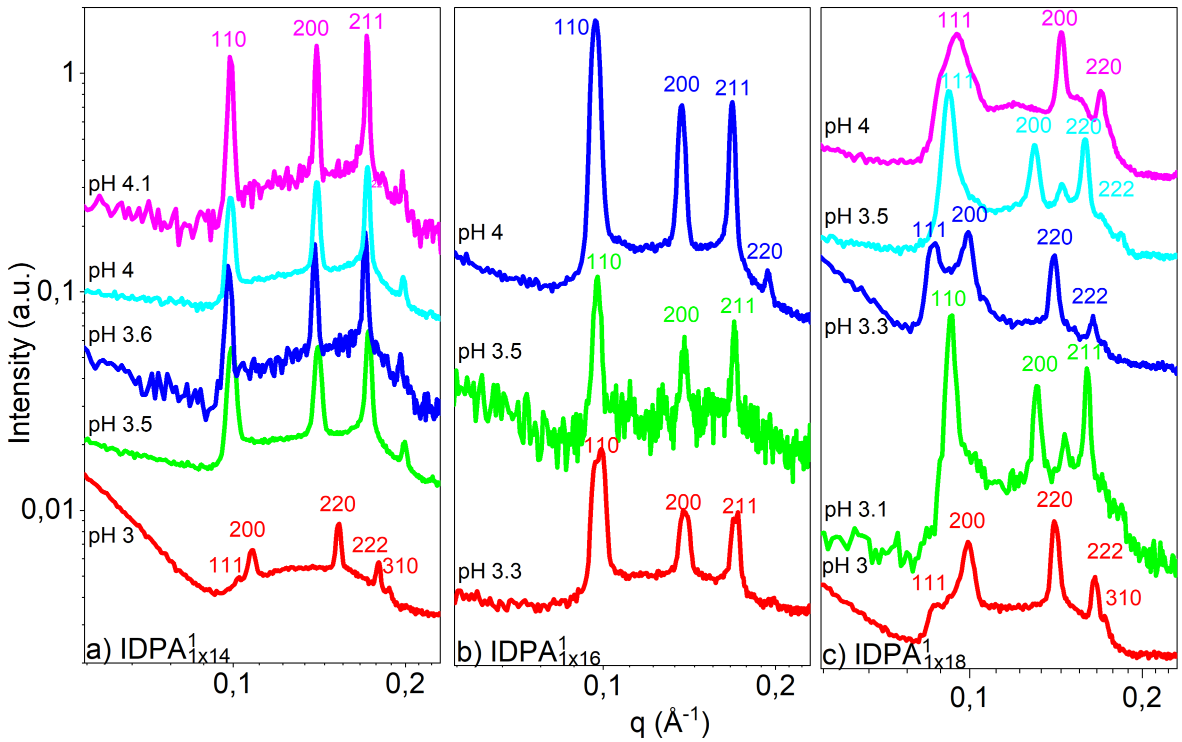


Figure S.12: **Scattering Signal for single tailed IDPAs** a) IDPA$^1_{1x14}$ b) IDPA$^1_{1x16}$ c) IDPA$^1_{1x18}$. Every IDPA was measured at different pH values, data was radially integrated and peak position was extracted using a Lorentian fit. Miller indices are shown for BCC and FCC lattices.

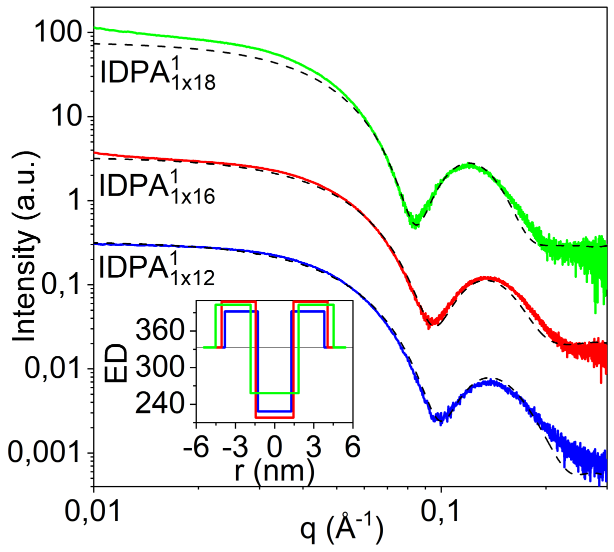


Figure S.13: **SAXS signal and fits for single tailed IDPAs** Dotted lines shows spherical form factor fit, Inset shows electron density profile for all three IDPAs. Data was fitted with X+[63] using a spherical form factor.

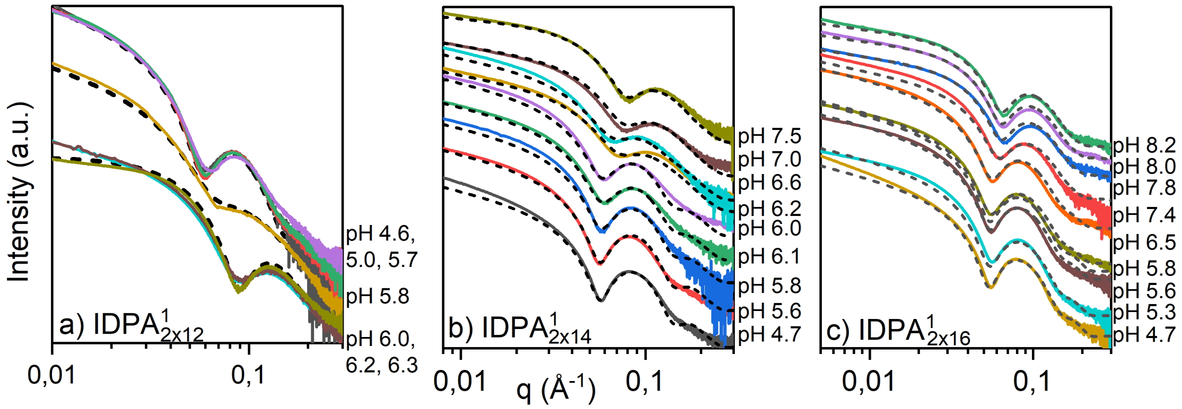


Figure S.14: **Sphere to Cylinder transition** Dotted lines show superposition of spherical and cylindrical form factors. Graphs are shown for a) IDPA$^1_{2\times12}$, b) IDPA$^1_{2\times14}$ and c) IDPA$^1_{2\times16}$.

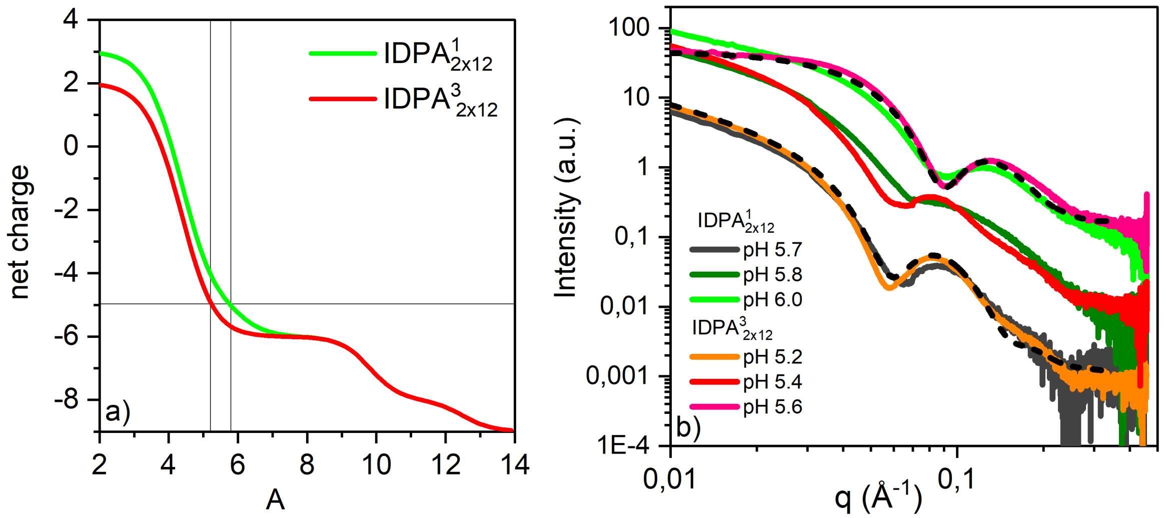


Figure S.15: **Spherical to worm-like micelle transition for IDPA$^1_{2\times12}$ and IDPA$^3_{2\times12}$** a) pH dependent charge projection predicts IDPA3 to transition at pH 5.2. Charge is calculated using Acetylation for N-terminus. b) Experimental SAXS data shows transition at pH 5.4.

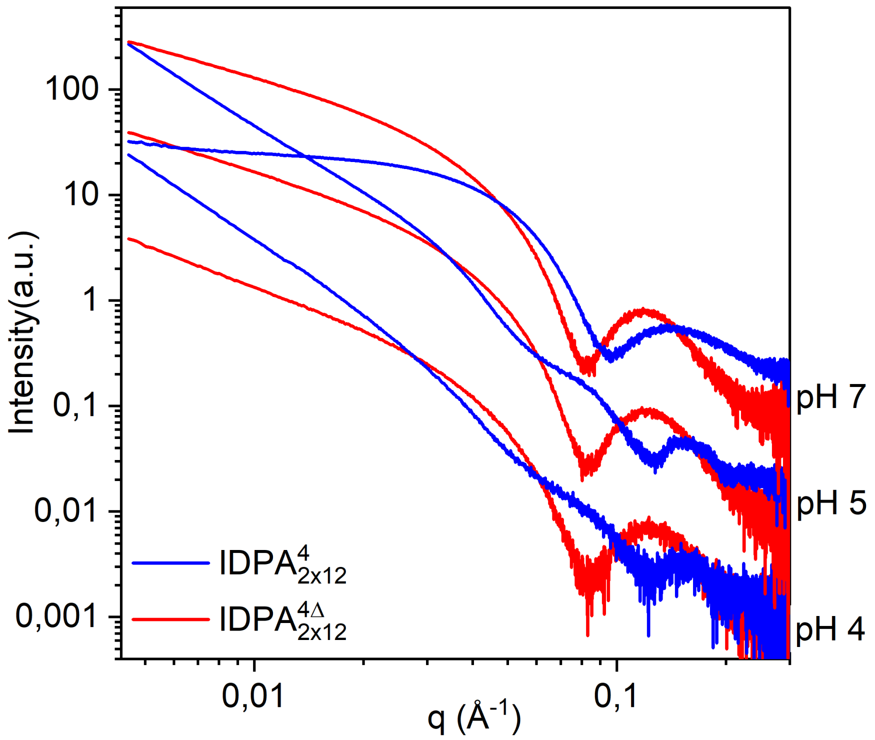


Figure S.16: **SAXS signal for IDPA$^{4}_{2\times12}$ and IDPA$^{4\Delta}_{2\times12}$** Cleaved IDPA$^{4\Delta}_{2\times12}$ remains at cylindrical form factor while IDPA$^{4}_{2\times12}$ undergoes phase transition. At physiological pH (pH 7) IDPA$^{4}_{2\times12}$ forms worm-like micelles. SAXS pattern for IDPA$^{4}_{2\times12}$ at pH 5 points towards the formation of polymer vesicles upon stretching of spherical micelles as shown by Takahashi et al.[61]

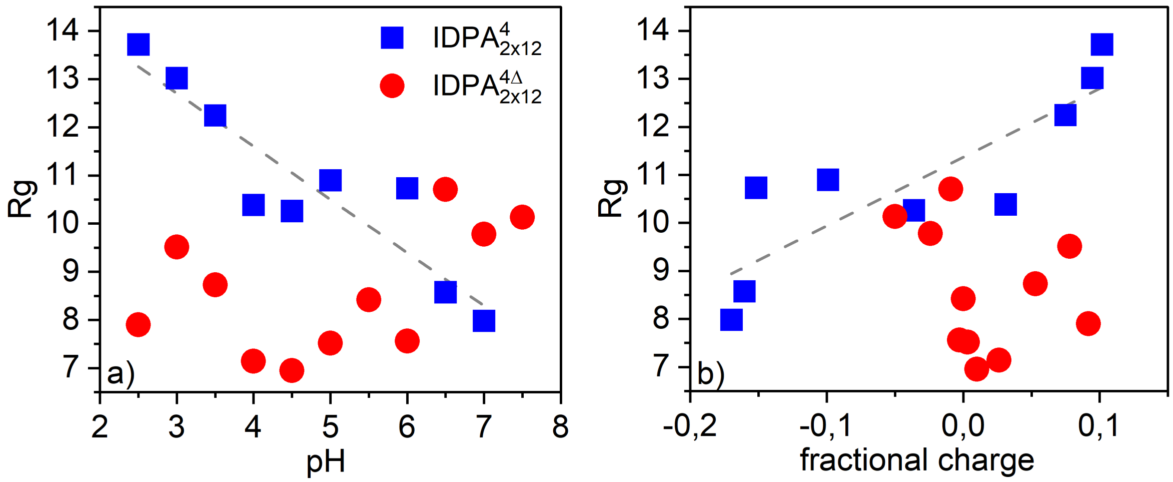


Figure S.17: **Peptides' scaling** a)$R_g$ versus pH b) $R_g$ versus fractional net charge. Line is a guide to the eye for tendency of IDP[4].

### SAXS Analysis

SAXS data was collected at various synchrotron sources (DESY, Hamburg, Germany Diamond light Source, Didcot, UK, Soleil, Paris France). The raw 2D Data was radially integrated using our in-house developed tool SAXSi.[46] We used X+ for fitting the core shell particels.[63] We used a spherical form factor to fit the micelles:

For spherical micelles we used the following form factor:

$$F_{sphere}(q,r) = \frac{3}{V_s} V_c \Big[\rho_c(r) - \rho_s(r)\Big] \frac{sin(qr_c) - qr\ cos(r_c)}{qr_c} + V_s \Big[\rho_s(r) - \rho_{solv}(r)\Big] \frac{sin(qr_s) - qr\ cos(r_s)}{qr_s} \tag{4}$$

Here $V_s$ is the volume of the whole particle, $V_c$ is the volume of the core, $r_s$ is the radius of the particle's shell, $r_c$ is the radius of the core, $\rho_c$ is the scattering length density of the core, $\rho_s$ is the scattering length density of the shell, $\rho_{solv}$, is the scattering length density of the solvent. $\rho_s$ was calculated by summing over amino acids' charges. Polydispersity for micellar sizes was added due to natural variations between biological samples.

For worm-like micelles we used a cylindrical form factor:

$$F_{cyl}(q,\alpha) = (\rho_c - \rho_s)V_c \frac{sin\left[\frac{1}{2}qL\,cos(\alpha)\right]}{\frac{1}{2}qL\,cos(\alpha)} \frac{J_1\left[qR\,sin(\alpha\right]}{qR\,sin(\alpha)} - (\rho_s - \rho_{solv})V_s \frac{sin\left[\frac{1}{2}qL + T\,cos(\alpha))\right]}{\frac{1}{2}qL + T\,cos(\alpha)} \frac{J_1\left[qR + T\,sin(\alpha)\right]}{qR + T\,sin(\alpha)} \tag{5}$$

Here $\alpha$ is the angle between the axis of the cylinder and $q$, $V_s$ is the volume of the whole particle, $V_c$ is the volume of the core, $L$ is the length of the core, $R$ is the radius of the core, $T$ is the thickness of the shell, $\rho_c$ is the scattering length density of the core, $\rho_s$ is the scattering length density of the shell, $\rho_{solv}$ is the scattering length density of the solvent. The outer radius of the shell is given by $R+T$ and the total length of the outer shell is given by $L+2T$. $J1$ is the first order Bessel function.

Both form factor were used to fit the data, Fig. S.18 shows how a sperical form factor captures the micelles, specifally in the low $q$ region.

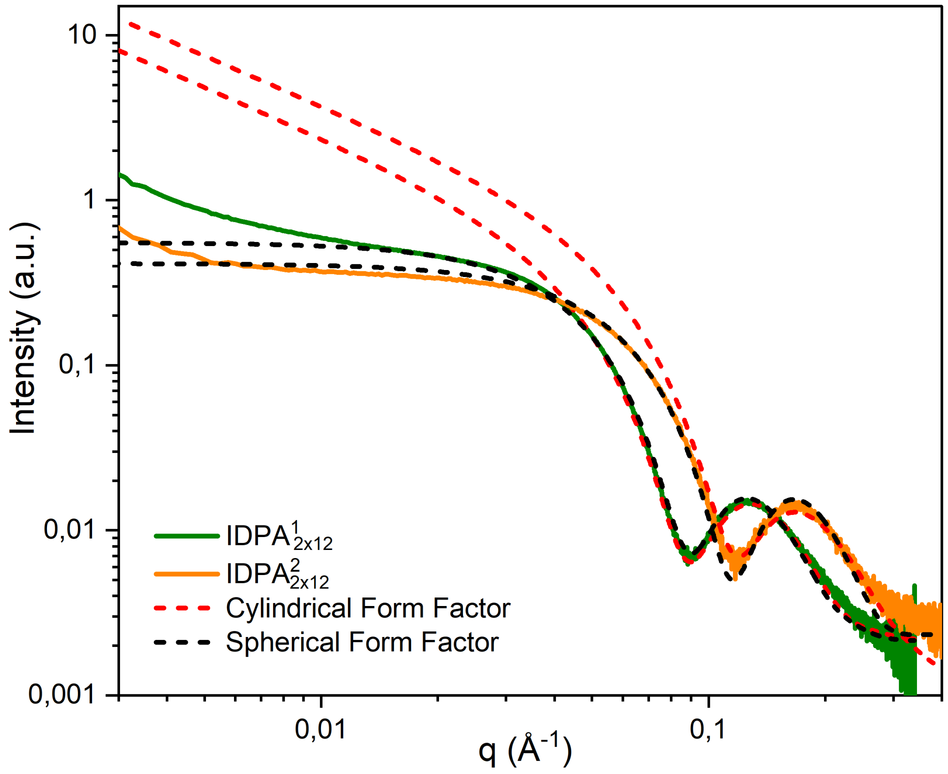


Figure S.18: **Spherical and cylindrical form factor for IDPA$^1_{2\times12}$ and IDPA$^2_{2\times12}$ at pH 6** Especially in for low $q$ values, a spherical form factor captures the data very well.
